# Performance of Sequential Markovian Coalescence Methods when Populations are Structured

**DOI:** 10.1101/2025.10.09.681379

**Authors:** Alba Nieto, Oscar Lao, Stefano Mona

**Affiliations:** Institut de Systématique, Evolution, Biodiversité (ISYEB), Muséum national d’Histoire naturelle, CNRS, Sorbonne Université, Université des Antilles, Paris, France; EPHE, PSL Research University, Paris, France; Group of Algorithms for Population Genomics, Department of Genetics, Institut de Biologia Evolutiva, IBE, (CSIC-Universitat Pompeu Fabra), Barcelona, Spain

**Keywords:** population structure, demographic inference, Sequential Markovian Coalescent, PSMC, SMC++, inverse instantaneous coalescent rate (IICR), inference bias

## Abstract

Sequentially Markovian Coalescent (SMC) methods are widely used to reconstruct demographic histories from genomic data, yet their accuracy in the presence of population structure has not been systematically evaluated. Here we assessed the performance of two popular SMC based algorithms (PSMC and SMC++) in retrieving the inverse instantaneous coalescent rate (IICR) simulated under two structured equilibrium scenarios, namely the finite island (FIM) and the two-dimensional stepping-stone (2D-SST). Looking backward in time, the IICR of a couple of lineages collected in a deme is characterized by a short period of constant value (the scattering phase) followed by an increase over time (the transition phase) up to a plateau (the collecting phase). Both algorithms recovered the IICR at the collecting phase but significantly deviated from the values expected during the transition phase under specific parameters’ combination. Specifically, the total error increased with the number of demes and the abruptness of the transition phase, which in turn is related to the extent of between demes connectivity. Moreover, each algorithm displayed a specific bias, corresponding backward in time to an artificial expansion either at the beginning of the collecting phase (PSMC) or of the transition phase (SMC++). Our results demonstrate that the observed systematic biases carry information about metapopulation dynamics, rather than being simple artefacts. Moreover, combining the results obtained from the two algorithms will increase our understanding of the historical demography of the metapopulations and help to put forward specific evolutionary scenarios to be tested with model-based algorithms using SFS or LD statistics.

## Introduction

With the rapid advancement of whole-genome sequencing, in-depth exploration of genetic variability has become a standard practice, even for non-model organisms. This revolution has positioned population genetics as a key discipline for unraveling species dynamics and providing crucial insights for conservation efforts (Ellegren, 2014).

Sequentially Markovian Coalescent (SMC)-based algorithms (McVean and Cardin, 2005; Wiuf and Hein, 1999), such as PSMC (Li and Durbin, 2011) and SMC++ (Terhorst et al., 2017), are state-of-the-art tools to recover population historical demography in extant and extinct species. PSMC and SMC++ use different summary statistics to compute the distribution of coalescence times of a couple of lineages (*T*_2_) along their genome, each with its strengths and limitations (Beichman et al., 2017; Mather et al., 2020; Peede et al., 2025; Sellinger et al., 2021). PSMC employs the distribution of heterozygote positions of a single diploid individual (or a couple of haploid lineages), which makes it especially valuable in the study of ancient samples or populations with scarce data (Bazzicalupo et al., 2022; Palkopoulou et al., 2015). SMC++, on the other hand, leverages the site frequency spectrum (SFS) from multiple individuals as well, increasing the accuracy of demographic inference in more recent times — a period where PSMC tends to underperform (Mather et al., 2020; Santiago et al., 2020; Schiffels and Durbin, 2014; Terhorst et al., 2017).

While often described as ‘model-free’ (Mather et al., 2020; Peede et al., 2025) because they do not assume a specific parametric function of population size change, these methods are still model-based in that they assume a panmictic population and neutral evolution under the coalescent with recombination. Typically, both algorithms estimate the (*T*_2_) distribution is then interpreted as the trajectory of changes in effective population size (*Ne*) through time (Li and Durbin, 2011; Terhorst et al., 2017). However, this interpretation is only valid under the assumption of panmixia, a condition rarely met in natural populations (Charlesworth et al., 2003; Moritz and Agudo, 2013). In reality, the expected distribution of *T*_2_ is influenced by all the underlying demographic processes at a time (Peede et al., 2025). Consequently, these trajectories are now more accurately referred to as the inverse of the instantaneous coalescent rate (IICR) (Chikhi et al., 2018; Mazet et al., 2016; Rodríguez et al., 2018). Under the presence of population structure, the expected IICR will reflect the structured coalescence (Chikhi et al., 2018; Herbots, 1994; Mazet et al., 2016). Typical structured models are the Finite Islands Models (FIM) (Wright, 1931), in which equal subpopulations called demes interchange individuals between all of them at the same time at a constant rate M, and the two-dimensions Stepping Stone (2D-SST) (Kimura and Weiss, 1964), where the subpopulations are only connected to the adjacent demes. In the latter, the deme position in the grid determines the level of isolation. As formulated in Wakeley (1999), the coalescent history of a sample of lineages from a single deme belonging to a metapopulation is typically divided, for mathematical tractability, into two distinct phases: the scattering phase and the collecting phase. The scattering phase refers to a period of rapid intra-deme coalescence predominating in recent times, with an expected coalescence rate of 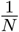, *N* being the diploid deme size. The collecting phase starts when all remaining non-coalesced lineages have migrated out of the deme, with each lineage being in a different deme. The rate of coalescence is lower in the collecting than in the scattering phase and this depends on the parameters of the structured model. In a finite island model, this rate increases with the value of migration rate and decreases with the within-deme *N* and with the number of demes. Backward in time, the two phases are bridged by a transition period (hereafter, transition phase) in which the coalescence rate gradually decreases, reaching a plateau which corresponds to the rate of the collecting phase.

Despite containing useful information on the metapopulation dynamics (Arredondo et al., 2021; Mazet et al., 2016; Wakeley, 1999), IICR trajectories inferred by SMC-based algorithms cannot inform on whether the population is structured or panmictic. For any structured model, there will always exist a panmictic model with variable effective population size producing exactly the same IICR.

SMC-based tools that infer the IICR, such as PSMC and SMC++, were developed and tested under the premises of a single panmictic population experiencing changes of *Ne* through time (Li and Durbin, 2011; Palamara et al., 2018; Santiago et al., 2020; Schiffels and Durbin, 2014). Yet, it remains unclear to what extent population structure introduces biases into the *Ne* trajectories retrieved by PSMC and SMC++.

In this study, we investigate whether population structure has an impact on the accuracy of PSMC and SMC++. Rather than focusing on departure from simulated Ne of the focal deme, we specifically investigate whether PSMC and SMC++, both assuming panmixia, can retrieve the expected IICR (*IICR*_*Sim*_) in the presence of population structure (*IICR*_*PSMC*_ and *IICR*_*SMC*++_). To this end, we explored a range of demographic parameters under two structured population models: FIM and 2D-SST. In the case of the 2D-SST, we also evaluated the impact of two sampling schemes (i.e. sampling from a central deme or from a corner deme).

## Methods

### Obtaining the IICR

The IICR can be calculated given the distribution of coalescence times for pairs or lineages (*T*_2_) applying Equation 1 from (Mazet et al., 2016), where 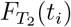 is the cumulative distribution of coalescent times and 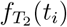 is the estimated density of coalescent events around *t*_*i*_.

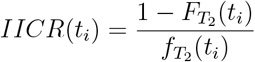

This distribution of *T*_2_ can be obtained from: (i) the theoretical prediction of the coalescent theory under specific (see below) demographic scenarios (*IICR*_*T*_ ) (Arredondo et al., 2021), (ii) by simulating a large number of independent *T*_2_ under any demographic scenario (*IICR*_*Sim*_), and (iii) by applying SMC approaches to genomic data (*IICR*_*Inf*_ ), based on the markovian coalescent approximation first proposed by (McVean and Cardin, 2005; Wiuf and Hein, 1999). The *IICR*_*T*_ can be calculated for any panmictic models and for specific structured demographic models at the equilibrium (Rodríguez et al., 2018), such as the FIM or 2D-SST. In such cases, the distribution of *T*_2_ can be mathematically derived under the non-stationary structured coalescence (NSSC) originally proposed by (Herbots, 1994) and further developed by (Rodríguez et al., 2018).

To obtain the inferred IICR (*IICR*_*Inf*_ ) we applied PSMC (*IICR*_*PSMC*_) and SMC++ (*IICR*_*SMC*++_) on simulated sequences. We averaged replicates to obtain one IICR trajectory per algorithm. Full simulation settings, parameter grids, and algorithm configurations are provided in the Supplementary Materials.

### Simulation Study

In this work we evaluated the performance of PSMC and SMC++ to retrieve the IICR when the lineages are sampled from a subpopulation (hereafter, a deme) belonging to a structured population (hereafter, a metapopulation). More specifically, we compared the *IICR*_*Sim*_ derived from the distribution of *T*_2_ simulated with *fastsimcoal*2 (Excoffier et al., 2013, 2021) to the *IICR*_*Inf*_ inferred by the two SMC-based algorithms. We considered two structured scenarios: FIM and 2D-SST. In FIM, the metapopulation is subdivided into *d* demes, each with a diploid population size *N* . All demes are interconnected, with 4*Nm* migrants exchanged per generation between a pair of demes, with *m* being the individual migration probability. The scaled migration rate (*M* ) is then calculated as *M* = 4*Nm*(*d−*1), representing the total number of emigrants from a deme per generation. All demes are equivalent in the FIM scenario, i.e., there is no spatial variability.

In the 2D-SST, the metapopulation is divided into d demes of equal diploid size (*N* ) arranged in a two-dimensional grid. Migration occurs only between geographically adjacent demes, resulting in spatially variable connectivity. The scaled migration rates equal to *M* = 4*Nm ×* 4 for central demes, as they exchange migrants with the four neighboring demes only. Conversely, it equals *M* = 4*Nm ×* 2 in corner demes, as they can only exchange with two adjacent demes. This spatial arrangement leads to variation in demes’ connectivity depending on the sampling location.

Specific parameters values for each demographic scenario were chosen in order to reproduce:

i) genetic diversity ranging between 1 *×* 10^*−*4^ and 1 *×* 10^*−*3^ per site, as measured by Tajima’s *θ* estimator (pairwise distance *θ*_*π*_); ii) *Fst* ranging between 0.01 and 0.3, to mimic differentiation values observed in natural species (Cho et al., 2013; Corbett-Detig et al., 2015; Dobrynin et al., 2015; Fischer et al., 2006; Lesturgie et al., 2022; Li et al., 2008; Omote et al., 2015; Prado-Martinez et al., 2013).

In FIM, we simulated 100 demographic models (group of models A) by generating all the possible demographic parameter combinations from a grid of values: *M* = [1, 2.5, 5, 15, 50], *d* = [5, 10, 20, 50] and diploid metapopulation size *Nd* = [2000, 10000, 20000, 35000, 50000]. In 2D-SST we considered square grids. We simulated 48 scenarios (group of models B), with demographic parameter combinations extracted from a grid of values of *M* (*M* = [1, 2, 4, 10]), *d* (*d* = [25, 49, 81, 121]) and diploid metapopulation size *Nd* (*Nd* = [5000, 10000, 20000]). To account for the spatial structure of 2D-SST, we considered two sampling schemes: the central deme of the grid of populations, or the corner deme (i.e. representative of the most and the least connected demes of the metapopulation, respectively). From each simulation we sampled 20 haploids –10 diploid individuals– from the same deme in each simulation, considering either the corner or the central deme in the 2D-SST and a random deme of the metapopulation in the case of FIM.

Finally, we compared both *IICR*_*Sim*_ and *IICR*_*Inf*_ between 2D-SST and FIM by fixing the same demographic values for each of the two models. Specifically, we generated 45 pairs of FIM and 2D-SST scenarios (group of models C) in which parameters were extracted from the grid: *M* (*M* = [2, 4, 8, 16, 40]), *d* (*d* = [25, 81, 121]) and diploid deme size *N* (*N* = [100, 150, 200]), with *M* referring to the central deme.

### Expected *IICR*_*Sim*_ analyses

#### Self-Organizing Maps (SOMs)

To visualize the relationship between the shape of *IICR*_*Sim*_ and the underlying demographic parameters, we reduced the dimensionality of the *IICR*_*Sim*_ shape using Self-Organizing Maps (SOMs) (Kohonen, 1982), employing the library *minisom* in *python* (Vettigli, 2018). SOMs is a type of artificial neural network architecture that maps high-dimensional input data to a grid of “neurons”, where each neuron represents a prototype vector. This mapping retains the structure and relationships in the data, with similar data points mapped close to each other. We choose a grid of four by four neurons after manually testing different configurations. We then examined the distribution of one demographic parameter at a time coloring the neurons based on the value of the demographic parameter for each *IICR*_*Sim*_ assigned to them. Additionally, we overlaid the shape of the most representative *IICR*_*Sim*_ (the “winner” *IICR*_*Sim*_) above each neuron.

To analyze the effect of the number of demes (*d*) on the *IICR*_*Sim*_ shape, we excluded scenarios with high connectivity (*M* = 50) in FIM (group of models A), which are nearly panmictic. In these cases, there is no difference in IICR values over time, regardless of the number of demes (Mazet et al., 2016; Wakeley, 1998, 1999). Such scenarios would hinder the proper functioning of the SOM.

#### Analysis of the transition phase

To assess the dynamics of the switch from the scattering to the collecting phase, we divided each simulated trajectory into 20 contiguous windows and computed the mean *IICR*_*Sim*_ value and corresponding mean time within each window. We then calculated slopes between all possible pairs of window means and recorded the maximum slope observed, which is an indicator of the abruptness of the transition phase. We applied the Wilcoxon paired ranks test to compare the maximum slope computed between the central and corner demes in the 2D-SST model (group B), as well as between pairs of models in the FIM and 2D-SST (group C) with equivalent demographic parameters.

We further evaluated whether the abruptness of the transition depends on the deme connectivity of the focal deme. In the FIM model this equals the number of demes minus one, whereas in the 2D-SST model it corresponds to the number of neighboring demes in the grid (two for corner demes, and four for central demes). We fitted a generalized linear model with the model type and sampling type as a categorical factor, thus testing the effect of deme connectivity on the slope independently of systematic differences between FIM and 2D-SST (corner and central demes).

### Quantification of the error of the SMC-based inference tools in structured populations

For a given demographic model and demographic parameters, we calculated the root of the mean squared error (*RMSE*) between the *IICR*_*Sim*_ and the IICR inferred by PSMC (*IICR*_*PSMC*_) and SMC++ (*IICR*_*SMC*++_) (See further details in Supplementary Material). We further built linear models relating the logarithm of the *RMSE* to each demographic parameter (N, M, d) to investigate their relative contribution to the total error. Using a Wilcoxon paired ranks test, we compared the *RMSE* values across i) FIM and 2D-SST models with equivalent demographic parameters (group of models C) and ii) different sampling strategies (central vs. corner deme) within the 2D-SST models (group of models B). Finally, to further evaluate the differences in performance of PSMC and SMC++ through time, we examined the distribution of squared errors, calculated as (*IICR*_*Sim*_-*IICR*_*Inf*_ )2 at each time point of vector v. We then assessed the uniformity of these errors using the Kolmogorov-Smirnov (K-S) test, which provides insights into the error of the inference over the temporal frame considered. Moreover, we evaluated the relative differences in error between both algorithms as:

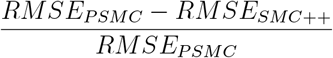

#### Bias of the inferences by PSMC in structured scenarios

When applying PSMC to our simulated datasets, we observed a recurrent overestimation of the *IICR*_*Inf*_ in the mid-ancient past (see Results). Under the assumption of panmixia, this overestimation would be interpreted as a past demographic expansion followed by a contraction. In equilibrium models, the maximum expected IICR corresponds to the coalescence rate during the collecting phase, which we approximated as the mean *IICR*_*PSMC*_ over the 10 most ancient time points. The intensity of the overestimation was computed as in:

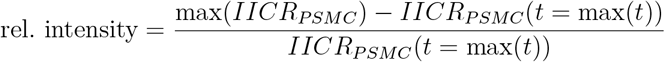

An artificial overestimation of the IICR was considered present when max(*IICR*_*PSMC*_) was at least 20% greater than the *IICR*_*PSMC*_ at the collecting phase, representing a representing a relative difference equal or larger than 1.2. This difference was considered large enough to suggest that the observed reconstruction was not simply due to the stochastic variation of the inferential process, but rather a systematic artifact.

We first correlated the relative intensity of the artificial expansion with the *RMSE*_*PSMC*_ to evaluate the impact of the artifact in the general error measurement. To further investigate the consequences of this bias, we divided the time series in three time transects: before, during and after the artificial expansion. Forward in time, we defined the start of the expansion as the point at which *IICR*_*PSMC*_ began to increase with respect to the plateau reached in the collecting phase. The end of the expansion was defined as the time when the *IICR*_*PSMC*_ reached its maximum.

We calculated the *MSE*_*PSMC*_ for each of these three times transects, and calculated the percentage of the overall *MSE*_*PSMC*_ corresponding to each time transect. We also correlated the *MSE*_*PSMC*_ during the artificial expansion with the *MSE*_*PSMC*_ after the artificial expansion.

We applied a Wilcoxon paired ranks test to compare the artificial expansion between: 1) sampling sites (corner vs central deme) in a 2D-SST (group B); and 2) pairs of FIM, and 2D-SST with equivalent demographic parameters (group C). Additionally, we modeled the probability of finding the artificial expansion (*rel*.*intensity >* 1.2) as a function of the parameters of the model through a multiple logistic regression (group of models A and B). We tested all combinations of parameters (N, m, and d) against the null model, which includes only the intercept, and selected the best model using the Akaike Information Criterion (AIC).

#### Biases of the inferences by SMC++ in structured scenarios

We observed a recurrent overestimation of the IICR by SMC++ (*IICR*_*SMC*++_) in FIM but not in 2D-SST scenarios in recent times. This artificial pattern mimics a recent population recovery in which the IICR, forward in time, starts to increase from its minimum value until the present time (note that in our study the most recent time considered is 50 generations). To quantify the intensity of this artificial recovery pattern, we calculated the difference between the minimum value of the *IICR*_*SMC*++_ and the value estimated in the most recent generation considered (corresponding to 50 generations, see Supplementary Materials) :

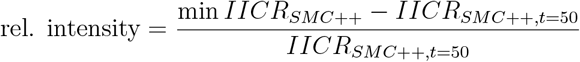

We defined the presence of an artificial recovery when the relative intensity exceeded 0.5, indicating a notable increase in the estimated IICR over the recent generations. To investigate the factors contributing to this bias, we conducted a multiple logistic regression analysis to model the probability of detecting a recent artificial expansion as a function of the demographic parameters of the model. We tested all combinations of parameters (*N, m*, and *d*) against a null model including only the intercept and identified the optimal model using the Akaike Information Criterion (AIC). This analysis resulted in a logistic regression model for the FIM simulations (group of models A). Additionally, we applied a logistic regression to explore the relationship between the probability of detecting this bias and the *Fst* values calculated from the simulated genomic data in FIM models.

To clarify the impacts of the recent artificial recovery in SMC++, we divided the time series into two intervals. Backward in time, the first time interval is defined from *v* = 50 generations to the onset of the expansion (defined as the minimum value of *IICR*_*SMC*++_ observed), and the second time interval starts from this time point to the last time point considered (see Supplementary Materials). For these two intervals, we calculated the *MSE*_*SMC*++_ and determined the percentage contribution of each of them to the total *MSE*_*SMC*++_ .

## Results

### The expected IICR under structured models

Consistently with previous findings (Mazet et al., 2016), *IICR*_*Sim*_ in our structured models tends to be lower in recent times compared to older times, with a transition phase bridging the scattering (recent times) and the collecting phase (older times). The specific trajectory of IICR variation depends on the combination of the three demographic parameters (*N, m*, and *d*) as well as the sampled deme in the case of 2D-SST scenarios (Figure 1). in two dimensions using a SOM (Supplementary Figure S1, S2). For both FIM and 2D-SST, the map reveals a structured gradient: the node in the upper left corner depicts the strongest difference between the maximum IICR value (reflecting ancient times in the collecting phase) and the minimum IICR value (reflecting recent times in the scattering phase), whereas the node in the lower right corner represents trajectories with the smallest difference between these two phases. The gradient in the IICR shape depends on the parameters of the underlying demographic models. In the case of FIM, gradients in the bidimensional space defined by SOM can be observed as a function of the migration rate (*M* ; Supplementary Figure S1A), the metapopulation size (*Nd*; Supplementary Figure S1B), and to a lesser extent the number of demes(*d*; Supplementary Figure S1C). For the 2D-SST model, *IICR*_*Sim*_ trajectories are similarly organized along gradients of *M* and *Nd* (Supplementary Figure S2).

**Figure 1.**
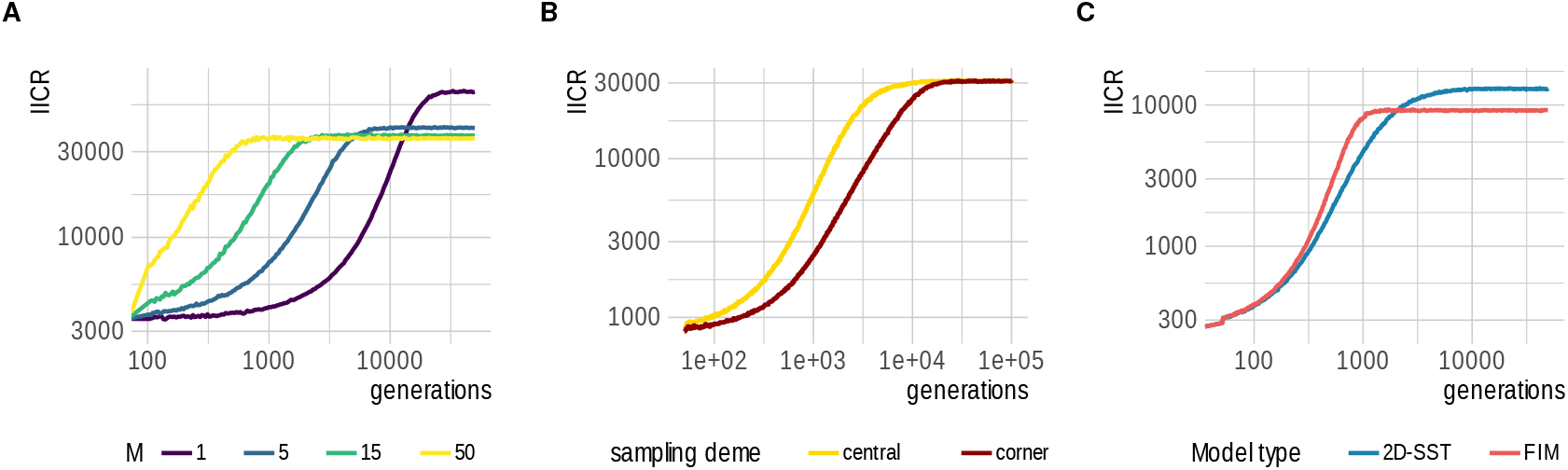
*IICR*_*Sim*_ changes depending on the parameters of the simulated model. Examples of recovered *IICR*_*Sim*_ from five (**A** and **C**) FIM and two (**B** and **C**) 2D-SST. Vertical axis represents the IICR value, and the horizontal axis the time measured in generations from the present to the past. The value of *IICR*_*Sim*_ surpasses *Nd* following the expected rate in the collecting phase in a structured scenario, which is proportional to *Nd* and inversely proportional to *M* . **A**. IICRS from three FIM models with fixed values metapopulation size of *Nd* = 35000 and *d* = 10. Scaled migration rate *M* value is 1 (purple), 5 (blue), 15 (green), and 50 (yellow). **B**. *IICR*_*Sim*_ from a 2D-SST model with *M* = 1, *d* = 25 and metapopulation size of *Nd* = 20000, sampled from a corner deme (red line) and from the central deme (yellow line). **C**. *IICR*_*Sim*_ from a pair of FIM (red line) and 2D-SST by sampling from central deme (blue line) models both with values of *M* = 2, *d* = 25 and deme size of *N* = 250.

We also compared the shapes of the *IICR*_*Sim*_ depending on demographic models. The abruptness of the transition phase was steeper in FIM than in the central deme (Wilcoxon’s paired rank test *p* − *value* = 3.06 *×* 10^*−*6^) (Figure 1C) and in the central deme than in corner deme of a 2D-SST scenario (Wilcoxon’s paired rank test *p value* = 5.62 × 10^*−*10^) (Figure 1B).

We further tested whether the abruptness of the transition is associated with the connectivity of the focal deme. A linear regression showed that the number of connected demes had a positive relation with abruptness (*β* = 0.025, *p − value* = 2.20 *×* 10^*−*16^), even when accounting for differences between demographic scenarios. This indicates that less isolated demes tend to experience a more abrupt transition, regardless of model type.

### Error and Biases in the inferred IICR by SMC-based methods Error in PSMC

To evaluate whether the demographic parameters of the model influence PSMC performance, we first examined the association between *RMSE*_*PSMC*_ and *M* (scaled migration rate), *d* (number of demes), and *N* (diploid deme size) fitting a linear model across both FIM and 2D-SST datasets.

In FIM models, *RMSE*_*PSMC*_ significantly increased with *d* (*β* = 1.19, *p*-value = 4.72 *×* 10^*−*13^) (see Figure 2,3), but decreased as deme size (*N* ) increased (*β* = *−* 0.70, *p*-value = 2.96 *×* 10^*−*3^). The error also increased with *M* when the number of demes was small (*d <* 50; *β* = 0.41, *p*-value = 2.35 *×* 10^*−*6^) (Figure Figure 2,3). Interestingly, the relationship between *RMSE*_*PSMC*_ and the value of *M* was inverted when the number of demes was large (*d* = 50; *β* = *−* 0.76, *p*-value = 1.0 *×* 10^*−*4^) (Figure Figure 2,3).

**Figure 2.**
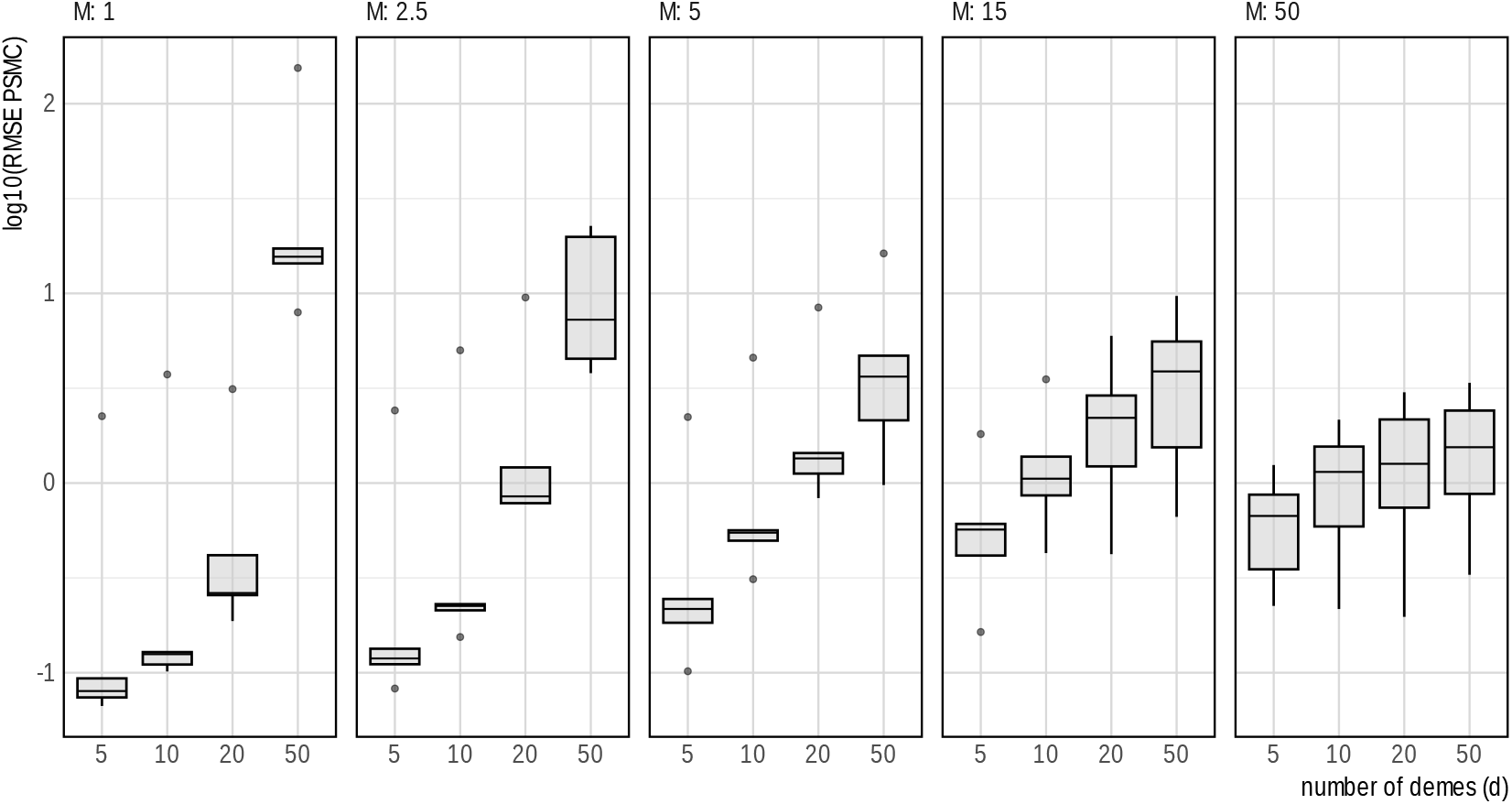
*IICR*_*PSMC*_ overall error in FIM. Panels from left to right correspond to values of scaled migration rate (*M* ) of 1, 2.5, 5, 15, and 50. Vertical axis represents the *RMSE*_*PSMC*_ in *log*10 scale and the horizontal axis represents the values of the number of demes (*d*).

**Figure 3.**
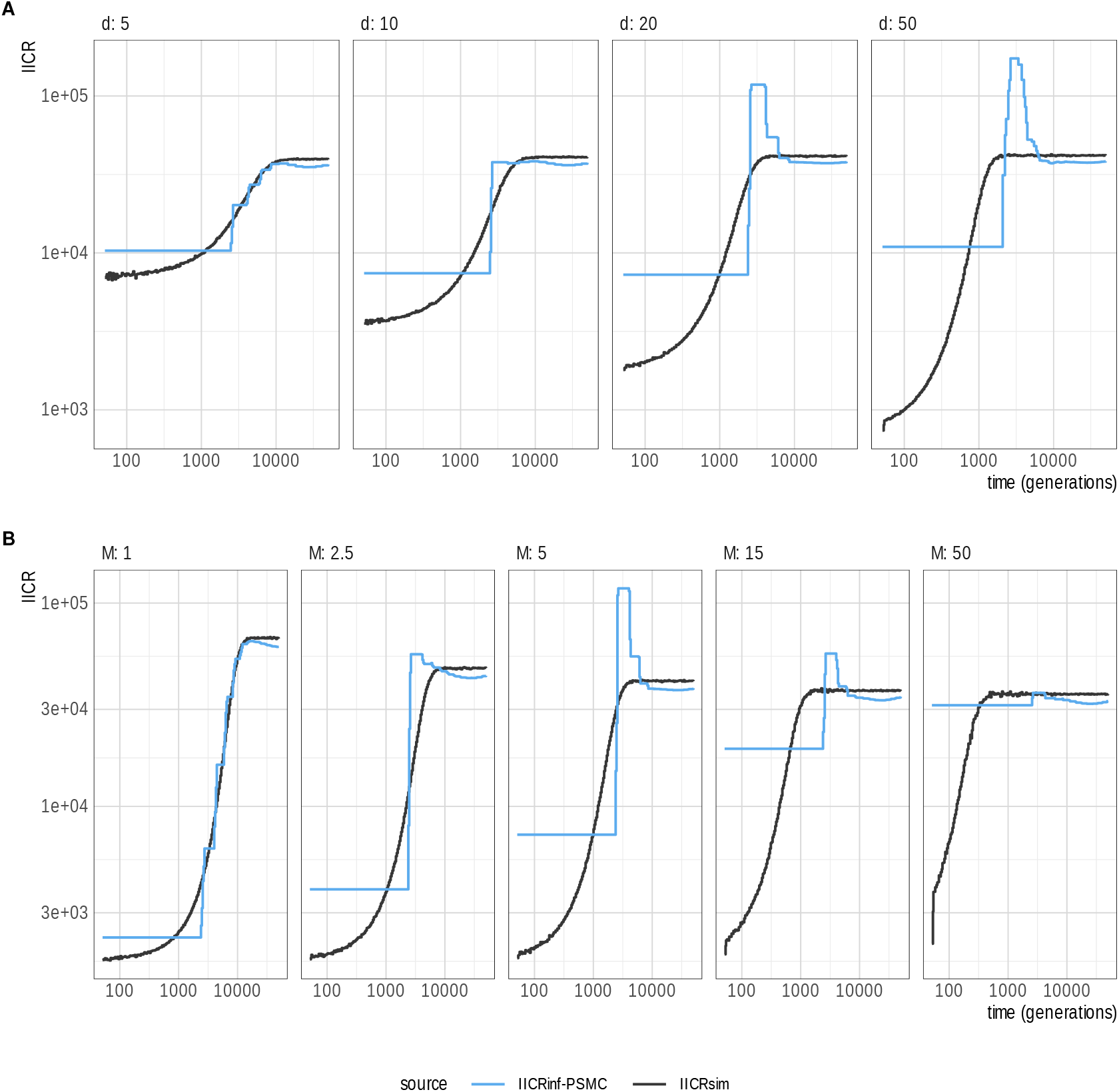
Comparison between *IICR*_*PSMC*_ and *IICR*_*Sim*_ in FIM models. Vertical axis represents the IICR value and horizontal axis the time in generations (from recent to ancient times). Simulate parameter values are *Nd* = 35000 and A. *M* = 5. Value of *d* changes from 5 to 50 from left to right panel. B. *d* = 20. Value of *M* changes from 1 to 50 from left to right panel. Blue lines: *IICR*_*PSMC*_; black lines: *IICR*_*Sim*_.

In 2D-SST models, we estimated the *RMSE*_*PSMC*_ values for central and corner demes given the different level of connectivity for the two sampling sites. *RMSE*_*PSMC*_ increased according to d in the corner demes (*β* = 0.41, *p − value* = 2.35 *×* 10^*−*3^) and in central demes (*β* = 0.31, *p − value* = 1.77 *×* 10^*−*2^) (Supplementary Figure S3).

We further compared *RMSE*_*PSMC*_ between FIM and 2D-SST models (considering the central deme only) with identical demographic values for *M, d*, and *Nd* (group C). Our analyses revealed that the *IICR*_*PSMC*_ was closer to the *IICR*_*Sim*_ in 2D-SST scenarios than in FIM (Wilcoxon’s paired rank test *p − value* = 1.4 *×* 10^*−*6^) (Figure 4A). On average, PSMC error was 1.56 units higher in FIM than in the equivalent 2D-SST models, (*CI*95% = [0.617, 2.497]).

**Figure 4.**
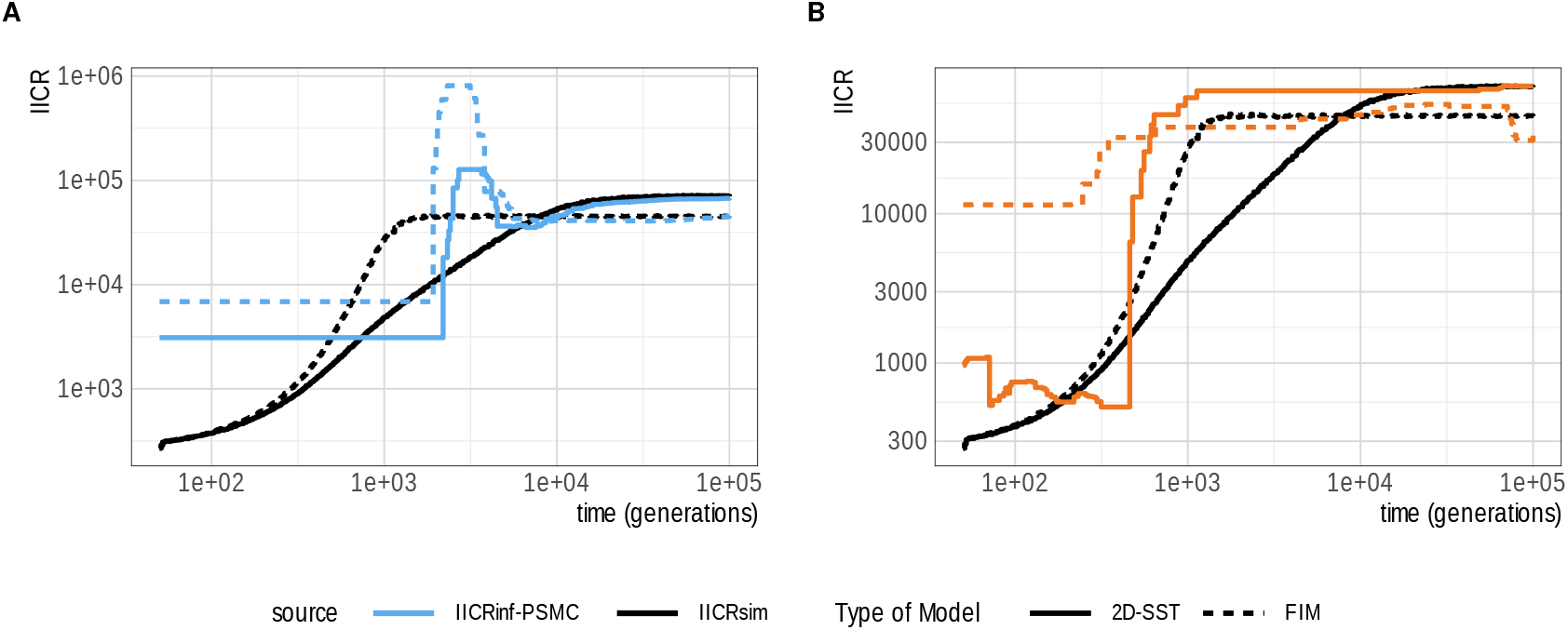
Comparison between *IICR*_*PSMC*_ (panel A) or *IICR*_*SMC*++_ (panel B) and *IICR*_*Sim*_ as a function of model type, FIM and 2D-SST. Vertical axis: IICR; horizontal axis: time in generations (from recent to ancient times). Simulated parameters are: *M* = 2, *d* = 121, *N* = 250. Black lines: *IICR*_*Sim*_ ; full lines: 2D-SST; dotted lines: FIM. **A**. Blue lines: *IICR*_*PSMC*_; **B**. Orange lines: *IICR*_*SMC*++_.

To further assess the impact of sampling, we compared *RMSE*_*PSMC*_ values for the *IICR*_*PSMC*_ computed from corner and central demes in the 2D-SST model. We observed a decline in PSMC performance in the central deme relative to the corner deme (Wilcoxon’s paired rank test *p − value* = 1.3 *×* 10^*−*9^) (Figure 5A). On average, the error of PSMC inferences is 0.71 units larger in the central deme than in the corner deme (*CI*95% = [0.50, 0.93]). Moreover, the difference in the error between central and corner demes showed a strong negative correlation with the migration rate *M* (*R*(*pearson*) = *−* 0.63, *CI*95% = [*−* 0.77, *−* 0.42]) (Supplementary Figure S3).

**Figure 5.**
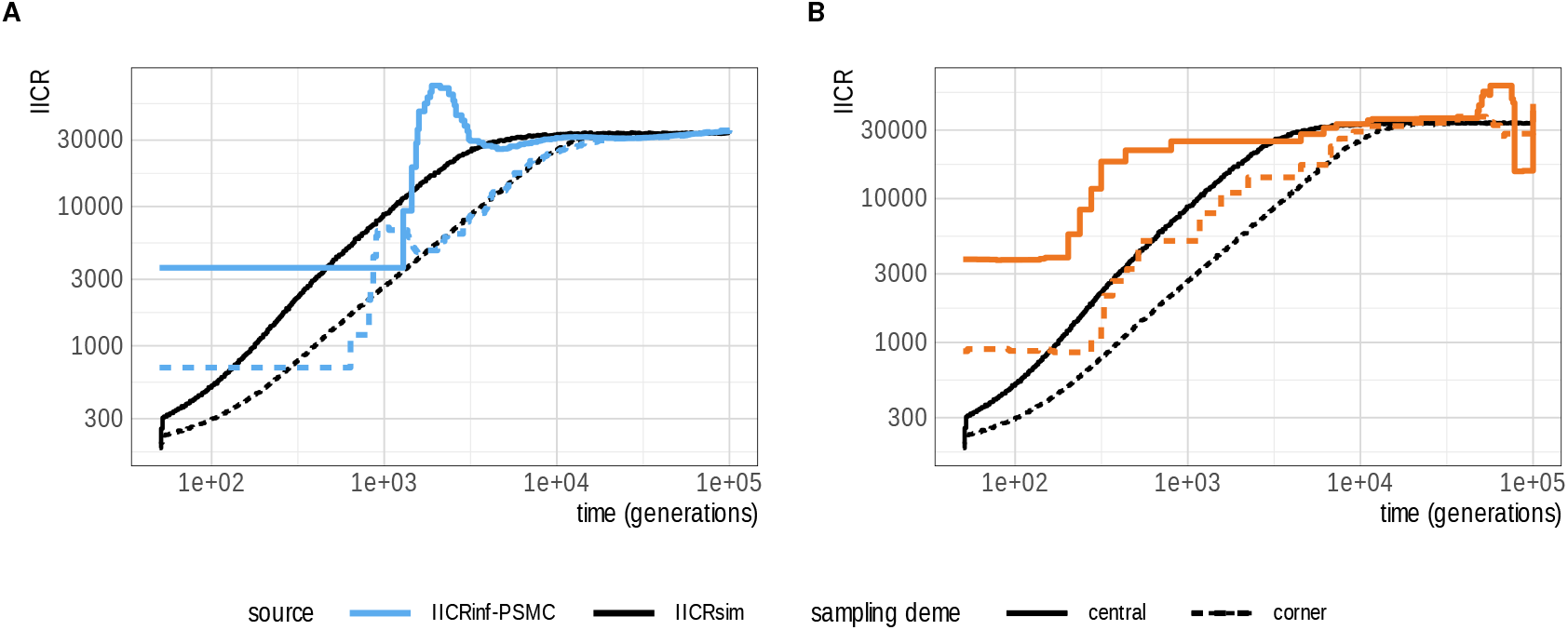
Comparison between *IICR*_*PSMC*_ (panel A) or *IICR*_*SMC*++_ (panel B) and *IICR*_*Sim*_ as a function sampling scheme in 2D-SST. Vertical axis: IICR; horizontal axis: time in generations (from recent to ancient times). Simulated parameters are: *M* = 2, *Nd* = 20000, *d* = 121. Black lines: *IICR*_*Sim*_ ; full lines: central deme; dotted lines: corner deme. **A**. Blue lines: *IICR*_*PSMC*_; **B**. Orange lines: *IICR*_*SMC*++_.

**Figure 6.**
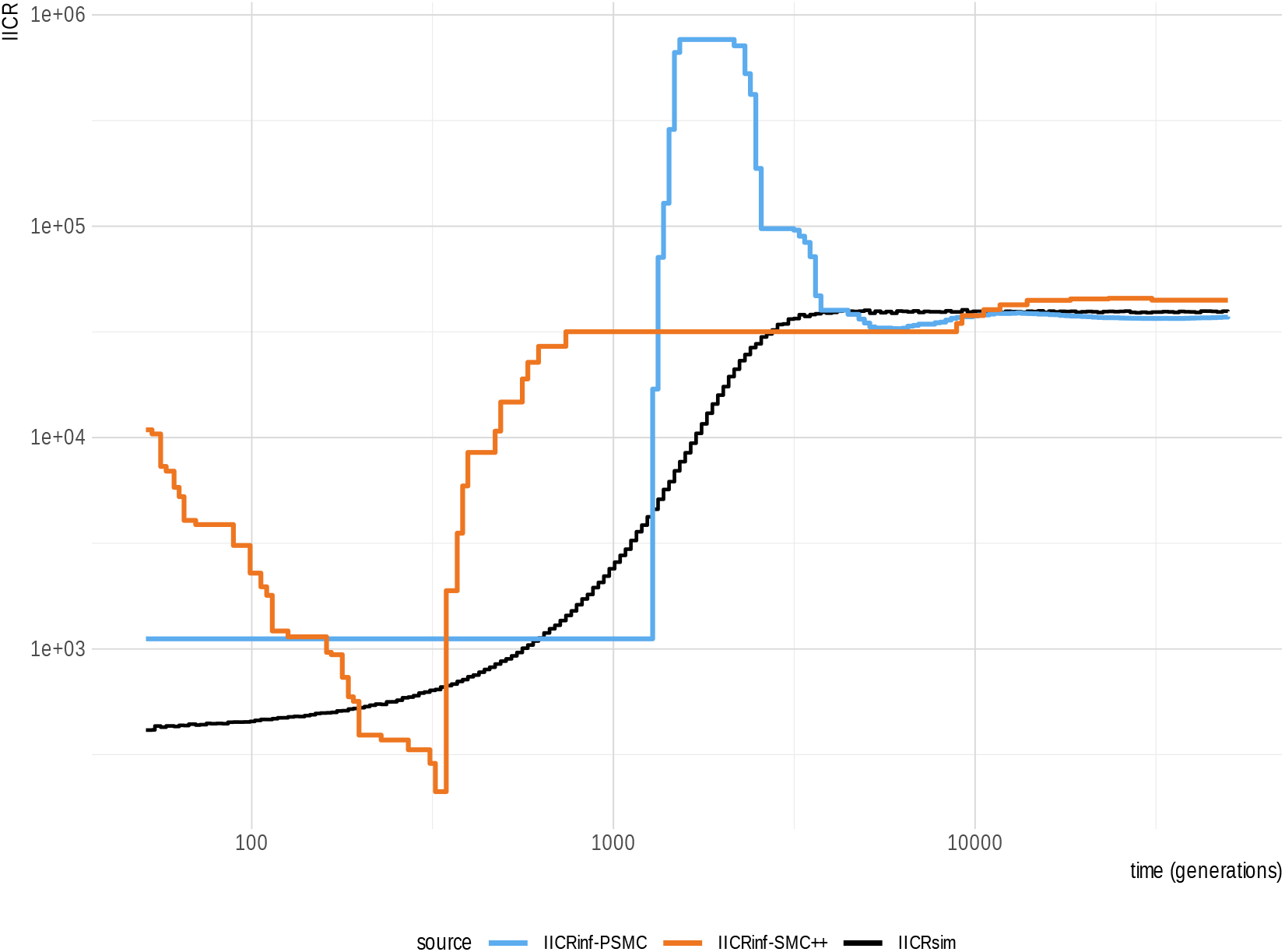
Systematic biases of *IICR*_*PSMC*_ and *IICR*_*SMC*++_ in FIM. Vertical axis represents the IICR value and horizontal axis the time measured in generations. Black line: *IICR*_*Sim*_; orange line: *IICR*_*SMC*++_; blue line: *IICR*_*PSMC*_. Simulated parameter: *d* = 50, *Nd* = 20000 and *M* = 1.

Finally, linear regression modelling showed that *RMSE*_*PSMC*_ increases strongly with the abruptness of the transition phase (*β* = 8.25, *p − value* = 1.80 *×* 10^*−*7^), even when controlling for model type and sampling site. Since abruptness is itself determined by the connectivity of the focal deme, we also evaluated this relationship directly and found that *RMSE*_*PSMC*_ increases with the number of connected demes (*β* = 0.29, *p −value* = 6.60 *×* 10^*−*6^), once again after accounting for model type and sampling site. This result suggests that PSMC error increases in more connected demes, due to the difference in the abruptness of the *IICR*_*Sim*_.

#### PSMC error through time

Next, we evaluated the uniformity of the distribution of *MSE*_*PSMC*_ with a Kolmogorov-Smirnov Test (KS). *MSE*_*PSMC*_ is not uniformly distributed in time neither in FIM (*mean*(*p − value*(*KS*)) = 1.16 *×* 10^*−*27^) nor in 2D-SST (*mean*(*p − value*(*KS*)) = 4.06 *×* 10^*−*117^).

We then characterized the recurrent overestimation of the *IICR*_*Sim*_ by PSMC (See Supplementary materials for all the inferences, and Figures 3,5A, 4A, and 6) at the onset of the transition phase (looking forward in time). Our results indicate a strong correlation between the relative intensity of the peak of *IICR*_*PSMC*_ and the total *RMSE*_*PSMC*_ in those scenarios where the bias is present both in FIM (*R*(*pearson*) = 0.801, *p − value* = 4.45 *×* 10^*−*19^) and in 2D-SST models (*R*(*pearson*) = 0.57, *p − value* = 1.04 *×* 10^*−*9^). However, we observed that the peak intensity varies across models and sampling strategies. Larger peaks occur in FIM compared to 2D-SST (Wilcoxon’s paired rank test *p − value* = 7.64 *×* 10^*−*4^) (Figure 4A). Within 2D-SST, sampling from the central deme amplifies the bias compared to corner deme (Wilcoxon’s paired rank test *p − value* = 1.02 *×* 10^*−*4^) (Figure 5A).

Since the major difference in the *IICR*_*Sim*_ between FIM and 2D-SST (and sampling site in 2D-SST) is the abruptness of the transition phase, we next tested whether the presence and intensity of the bump are linked to this feature. Logistic regression showed that the probability of detecting a bump increases with the abruptness of the transition phase (*β* = 0.78, *p − value* = 0.02), even after accounting for model and sampling type. Likewise, linear regression indicated that the intensity of the bump also scales positively with abruptness (*β* = 11.85, *p − value* = 8.71 *×* 10^*−*6^). Given that the abruptness is determined by the number of connected demes to the focal deme, we further tested this relationship and found that both the probability of detecting the bump (*β* = 0.05, *p − value* = 3.54 *×* 10^*−*4^) and its intensity (*β* = 0.40, *p − value* = 1.58 *×* 10^*−*4^) increase with the number of connected demes, independently of model type. This shows that the occurrence and strength of the PSMC artifact are strongly linked to the connectivity —and thus the isolation— of the deme where sampling takes place. Additionally, we divided the overall *MSE*_*PSMC*_ into three time transects (before, after, and during the artificial peak) to evaluate the relative contribution of each interval to the total error. Notably, the error computed after the onset of the expansion (backward in time), in the most ancient part of the inference, represents a relatively small portion of the total *MSE*_*PSMC*_ in FIM (0.82%, *CI*95% = [0.27%, 1.38%]) and 2D-SST (0.24%, *CI*95% = [0.17%, 0.32%]).

On the contrary, the artificial expansion period itself accounts for the 23.44% (*CI*95% = [15.61%, 31.29%]) and the 10.64% (*CI*95% = [7.39%, 13.90%]) of the total error in FIM and in 2D-SST scenarios respectively. Finally, the error associated with the initial time interval (i.e. from time 0 to the collapse of the *IICR*_*PSMC*_ after the peak) accounted for 75.72% (*CI*95% = [67.9%, 83.5%]) and 89.1% (*CI*95% = [85.8%, 92.3%]) of the total error in the FIM and 2D-SST scenarios, respectively. We further observed that the error in this transect is strongly correlated with the relative intensity of the peak in FIM (*R*(*pearson*) = 0.76, *p − value* = 6.89 *×* 10^*−*8^) than in 2D-SST (*R*(*pearson*) = 0.28, *p − value* = 0.027). This finding suggests that the greater the intensity of the peak, the larger will be the overestimation of the IICR by PSMC in recent times.

#### Bias in PSMC

Further analyses showed a direct link between the timing of the artificial expansion and the total simulated metapopulation size. A linear regression of log_10_(timing) against log_10_(*Nd*) yielded a slope of 0.92 (*p*-value *<* 2.20 *×* 10^*−*16^) and an intercept of *−* 0.73 (*p*-value = 1.20 *×* 10^*−*12^) in FIM and a slope of 0.92 (*p*-value *<* 2.2 *×* 10^*−*16^) and an intercept of *−* 0.70 (*p*-value = 8.0 *×* 10^*−*10^) in 2D-SST. These slopes, being close to 1, indicate that increasing *Nd* by a given factor results in an almost identical factor increase in the timing of the expansion.

We next assessed the probability of observing such an artifact as a function of the demographic parameters used in the simulation by comparing different multiple logistic regression models. In FIM, the best model was *p*_artifact(PSMC)_ = *β*_0_ + *β*_1_*M* ^2^ + *β*_2_*M* + *β*_3_*d*. This model shows a direct linear relationship with *d* (*β*_3_ = 1.46, *p*-value = 1.31 *×* 10^*−*4^) and a quadratic relationship with *M*, where the probability of observing the artifact is highest for mild levels of migration (*M* ) and decreases for low or high connectivity (Supplementary Figure S5, Figure 3). The artifact is more intense at low *M* values (*R* = *−* 0.70, *p*-value = 1.98 *×* 10^*−*7^) and when *d* is high (*R* = 0.57, *p*-value = 6.72 *×* 10^*−*5^) (Supplementary Figure S5, Figure 3).

In 2D-SST, the artifact emerges when sampling from the central deme. The best model for predicting the probability of the artifact expansion is: *p*_*artifact*(*PSMC*)_ = *β*_0_ + *β*_1_*M* . In this model, the probability of observing the artifact is inversely proportional to *M* (*β*_1_ = *−*3.09, *p − value* = 3.03 *×* 10^*−*3^). Moreover, when the bias is present, its intensity will be higher in scenarios of low connectivity (*M* ) (*R* = *−*0.76, *p − value* = 1.77 *×* 10^*−*8^).

### Error in SMC++

We investigated whether SMC++ performance varied across the same demographic scenarios and sampling schemes previously analyzed with the PSMC. We first compared the *RMSE*_*SMC*++_ between inferences performed in the central and the corner deme in 2D-SST scenarios (Figure 5B). We observed a significantly higher error in the central deme (Wilcoxon’s paired rank test *p − value* = 0.009), with a mean difference between the *RMSE*_*SMC*++_ at the central deme compared to the corner deme of 0.34(*CI*95% = [0.02, 0.66]), which means that the error of SMC++ is 0.34 units larger in the central deme (Figure 5B). Additionally, when comparing scenarios with the same demographic parameters, error rates of *IICR*_*SMC*++_ were significantly greater in FIM than in the central deme of the 2D-SST (*p − value* = 0.004), with a mean difference of 1.16(*CI*95% = [0.30, 2.02]) (Figure 4B).

Linear regression showed that *RMSE*_*SMC*++_ increases with the abruptness (*β* = 1.58, *p − value* = 1.60 *×* 10^*−*13^), even after controlling for model type and sampling site. Since abruptness is determined by the number of connected demes, we also tested this relationship directly and found that *RMSE*_*SMC*++_ increases with deme connectivity (*β* = 0.06, *p − value* = 2.50 *×* 10^*−*11^).

Higher *RMSE*_*SMC*++_ is observed in structured scenarios with a large number of demes (*d*) (for FIM, *β* = 0.83, *p − value* = 2.20 *×* 10^*−*16^; in 2D-SST, *β* = 0.62, *p − value* = 4.63 *×* 10^*−*8^) (Figures 7, 8, Supplementary Figure S4) and for decreasing N in both 2D-SST (*β* = *−*1.04, *p − value* = 2.53 *×* 10^*−*10^) and FIM (*β* = *−* 0.42, *p − value* = 3.17 *×* 10^*−*3^) models.

**Figure 7.**
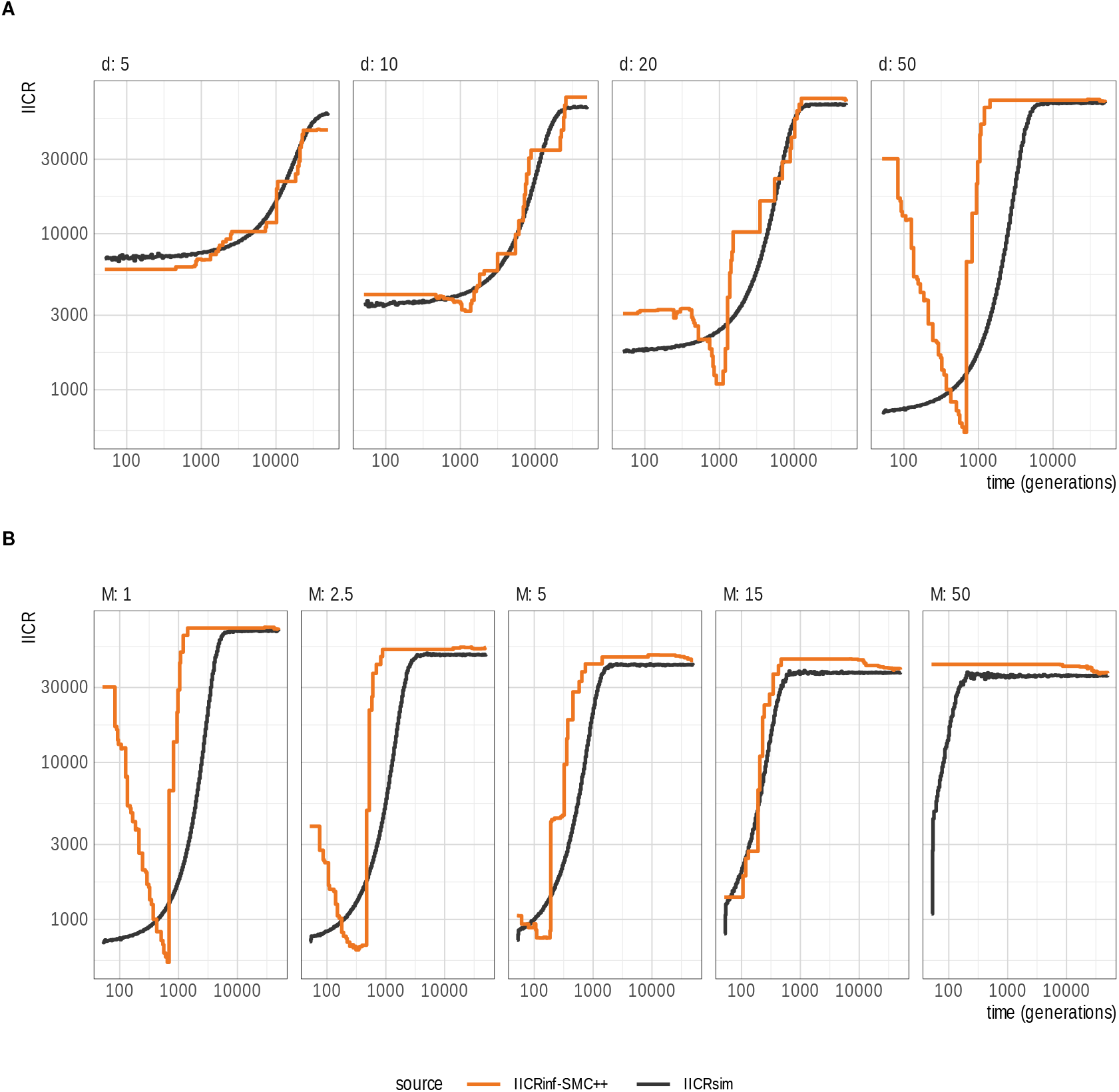
Comparison between *IICR*_*SMC*++_ and *IICR*_*Sim*_ in FIM models. Vertical axis: IICR values; horizontal axis: time in generations. Simulated parameters are *Nd* = 35000 haploids and **A**. *M* = 1. Value of d varies from 5 to 50 (from left to right). **B**. *d* = 50. *M* varies from 1 to 50 (from left to right). Orange lines: *IICR*_*SMC*++_, black lines: *IICR*_*Sim*_.

**Figure 8.**
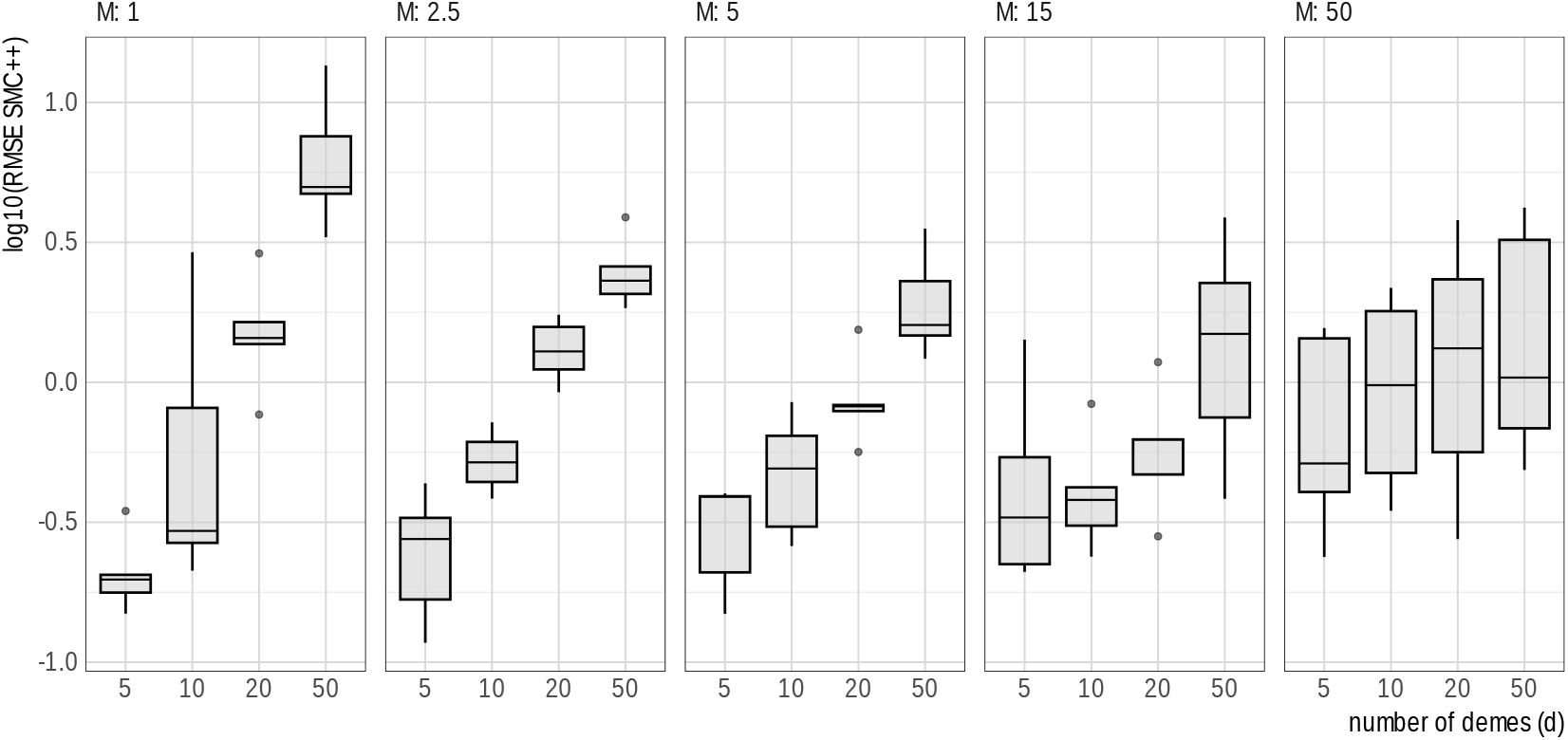
*IICR*_*SMC*++_ overall error in FIM. Panels from left to right correspond to values of scaled migration rate (*M* ) of 1, 2.5, 5, 15, and 50. Vertical axis represents the *RMSE*_*SMC*++_ in *log*10 scale and the horizontal axis represents the values of the number of demes (*d*)

### SMC++ error through time

Similarly to PSMC, we observed that the error of SMC++ was not uniformly distributed, showing a negative correlation between time and the *RMSE*_*SMC*++_ in both FIM (*R*(*pearson*) = *−* 0.47, *CI*95% = [*−* 0.37, *−* 0.56]) and 2D-SST (*R*(*pearson*) = *−* 0.48, *CI*95% = [*−* 0.55, *−* 0.41]).

Finally, we observed an overestimation of the *IICR*_*Sim*_ in recent times in FIM but not in 2D-SST (Figures 6, 7). When interpreting the IICR under the panmictic paradigm, this would mimic a population recovery or recent expansion. To further characterize which factors influence this artificial trend, we defined the expansion in mathematical terms by measuring the relative difference between the *IICR*_*SMC*++_ in the most recent generation considered (*t* = 50) and its value at the onset of the expansion, which in all cases corresponded to the minimum *IICR*_*SMC*++_. We consider that the SMC++ produces this artifact in a FIM simulation when the relative difference is bigger than 0.5. In these cases, the fraction of the total *RMSE*_*SMC*++_ due to the bias represents on average the 34.0%(*CI*95% = [12.76%, 55.25%]) of the total error in FIM models.

#### Bias in SMC++

The relationship between FIM demographic parameters and the probability of SMC++ producing this artificial expansion was analyzed by generating several multiple logistic regression models. The model considering all the three demographic parameters provided the best fit *p*_*artifact*(*SMC*++)_ = *β*_0_ + *β*_1_*M* + *β*_2_*d* + *β*_3_*N*, (*β*_0_ = *−* 4.96, *p − value* = 4.91 *×* 10^*−*4^). The parameter *d* exhibits strong positive association with the probability of finding the artificial expansion (*β*_2_ = *−* 2.24, *p − value* = 1.35 *×* 10^*−*2^), while *M* shows a strong negative relationship (*β*_1_ = *−* 2.95, *p − value* = 9.20 *×* 10^*−*3^). A weaker positive relationship is observed with *N* (*β*_3_ = 3.44, *p* − *value* = 6.32 *×* 10^*−*2^). These results imply that we expect to find the artificial expansion in scenarios with large number of demes (*d*), low connectivity (*M* ) and elevated deme size (*N* ) (Supplementary Figure S4). These conditions would correspond to scenarios with high differentiation between subpopulations. We further tested the association between the extent of population structure (as measured by *Fst*) and the probability of finding the artificial expansion. Consistently, a logistic regression *p*_*artifact*(*SMC*++)_ = *β*_0_ + *β*_1_*Fst* (*β*_0_ = *−* 4.11, *p − value* = 3.32 *×* 10^*−*4^) revealed that the probability of observing the artificial recent expansion is positively associated with the amount of population differentiation quantified by *Fst* (*β*_1_ = 1.81, *p − value* = 9.01 *×* 10^*−*3^). As an example, we found a probability of finding the artificial expansion higher than 50% in scenarios with *Fst* higher than 0.27 (Supplementary Figure S5).

#### Comparison of PSMC and SMC++ performance

We calculated the relative difference between *RMSE*_*PSMC*_ and *RMSE*_*SMC*++_ in order to test which algorithms performed better under the simulated scenarios.

In both FIM and 2D-SST scenarios, we found a significant correlation between the difference in *RMSE* between the two methods and the scaled migration rate (*M* ) (*R* = 0.64, *p − value* = 7.61^*−*7^), indicating that SMC++ increasingly outperforms PSMC as connectivity rises. In FIM models with many demes (*d* = 50), the performance of the two algorithms is similar. However, for smaller deme sizes (*d <* 50), differences emerge. At low migration rates (*M ≥* 2.5), PSMC performs better, with errors 55% smaller on average (*CI*95% = [44%, 66%]; Wilcoxon paired rank test, *p − value* = 5.96 *×* 10^*−*8^). In contrast, at higher migration rates (2.5 *< M ≥* 50), SMC++ outperforms PSMC, with errors 95% smaller on average (*CI*95% = [28%, 174%]; Wilcoxon paired rank test, *p − value* = 4.36 *×* 10^*−*3^). These findings demonstrate that PSMC excels at lower migration rates, while SMC++ becomes more effective as connectivity rises (Figure 9).

**Figure 9.**
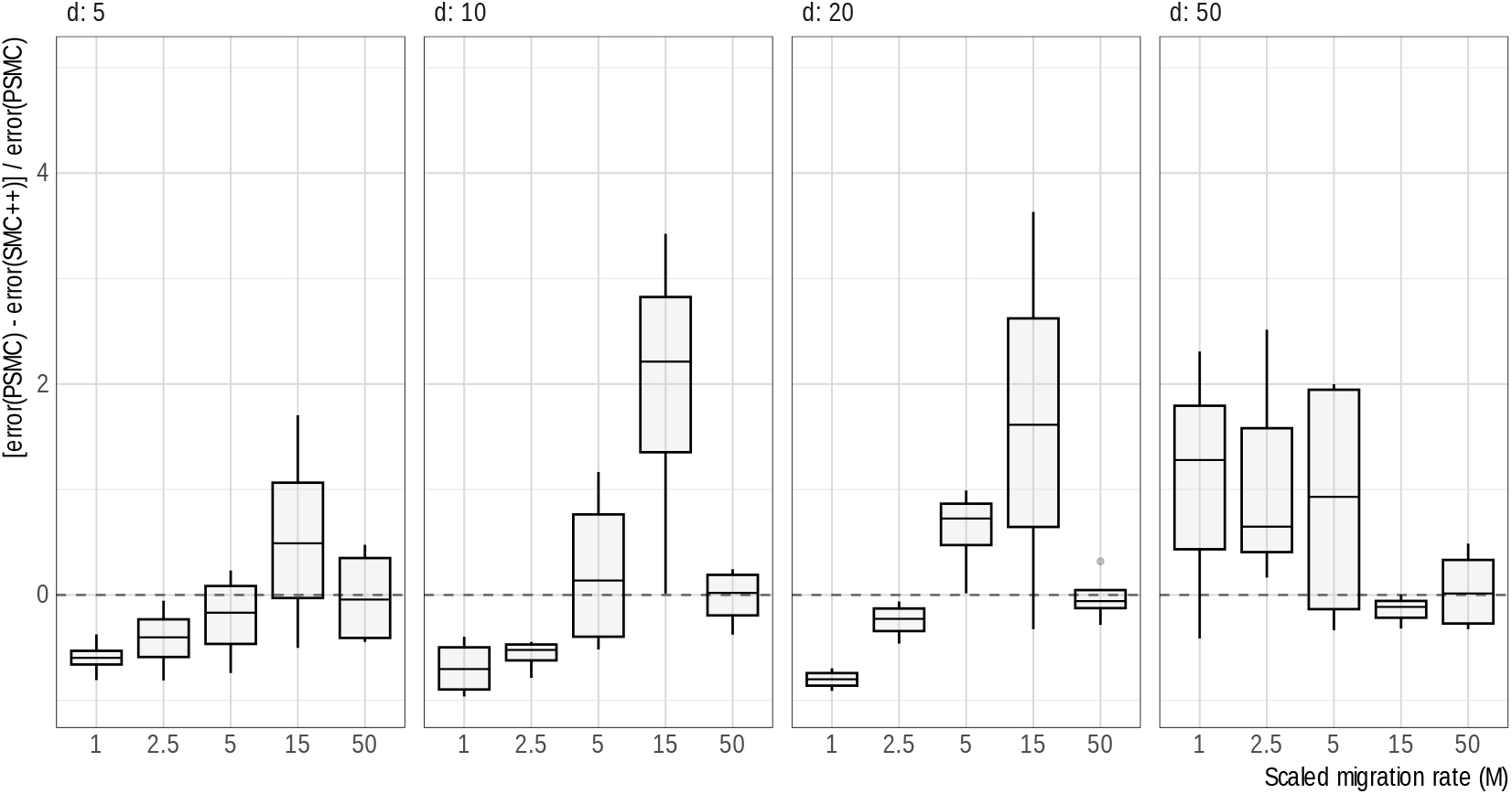
Difference in error between PSMC and SMC++ in FIM. Panels are arranged according to *d* = [5, 10, 20, 50] from left to right. Horizontal axis and color indicate the value of *M* and vertical axis the standardized difference in error between PSMC and SMC++. Negative values indicate a better performance of PSMC whereas positive values indicate a better performance of SMC++. When the difference is 0 (horizontal dashed line) the two softwares have an equivalent performance.

## Discussion

Methods based on the SMC theory reconstruct the history of coalescence events along the genome with the general underlying assumption of panmixia, with few exceptions (Cousins et al., 2025; Li and Durbin, 2011; Shchur et al., 2022; Wang et al., 2020). However, the distribution of coalescence events in structured populations also depends on the extent of population structure (and its variation through time), admixture events, population divergence, range contractions and expansions (Arredondo et al., 2021; Chikhi et al., 2018; Lesturgie et al., 2022; Mazet et al., 2016; Mona et al., 2014; Rodríguez et al., 2018).

Up to now, the ability of PSMC and SMC++ to reconstruct the true IICR has been only evaluated in panmictic (Li and Durbin, 2011; Sellinger et al., 2021; Terhorst et al., 2017) but not in structured models. To fill this gap, in this study evaluated how the two algorithms perform in reconstructing the IICR from diploid individuals sampled from a deme belonging to a metapopulation.

### Population Structure and IICR Dynamics

Our simulations show that the *IICR*_*Sim*_ was systematically lower in recent times, with a transition phase linking past and present coalescence rates, in agreement with previous studies (Chikhi et al., 2018). The scattering phase depends on within-deme *N*, while the collecting-phase plateau follows a Kingman process scaled by deme number and connectivity (Wakeley, 1999). The difference in IICR between ancient and recent times mostly depends on the migration rate *M* and number of demes *d*, with lower values of *M* and higher values of d producing larger differences between the maximum and minimum IICR. Indeed, reducing migration decreases the chance of two lineages ending up in the same deme to coalesce, increasing their MRCA (and the within-deme observed diversity) (Figure 1; Supplementary Figures S1, S2).

Conversely, at high migration the IICR resembles that of a panmictic population of size *Nd* (Figure 1). (Supplementary Figures S1, S2).While these results are consistent with earlier findings (see for example Mazet et al. (2016)), SOM analysis revealed that the IICR trajectory (the values in the scattering and collecting phase) strongly depends on the extent of population structure. More importantly, SOM analysis shows that the transition phase is not constant across simulations, but highly dependent on the demographic parameters of the model. Moreover, contrasting the IICR of different demes can inform on the spatial structure of the metapopulation, as more isolated demes will tend to display smoother transition phases.

Using linear modeling, we demonstrate that the steepness of the transition phase depends on the number of demes connected to the focal deme: the more isolated the focal deme is from the rest of the population, the less steep will be the shift from the scattering to the collecting phase (Figure 1). This implies that, when comparing scenarios with the same total number of demes and migration rate, we observed that the corner deme in 2D-SST scenarios has a smoother transition period than the central deme and any deme in a FIM model. The result can be understood intuitively: the less connected demes are, the longer it will take for the surviving lineages (i.e. those that did not coalesce within the deme) to escape from the sampled deme. The transition phase therefore harbours a mix of intra and inter-deme coalescences events before reaching the theoretical rate of the collecting phase.

### Biases in PSMC and SMC++ Inference

We assessed the performance of PSMC and SMC++ in structured models by comparing the inferred IICR to its expected value, generated by simulations, across a grid of FIM and 2D-SST models. Both algorithms can globally recover the IICR values corresponding to the collecting phase in all simulated scenarios for all combinations of demographic parameters. Conversely, the recent IICR, corresponding to the scattering phase, was not always well recovered (Figures 3, 4, 5, 6, 7. In both algorithms, we found that the error increased with the number of demes, which also corresponds to larger differences between the scattering and the collecting phase (Figures 2, 6). Moreover, we showed that the *RMSE* increased in both algorithms with the abruptness of the transition phase, which in turn depends on the number of demes connected to the focal deme. These results are consistent with previous findings (Sellinger et al., 2021), showing that SMC-based algorithms perform poorly in the presence of abrupt changes. Here we extended this conclusion by demonstrating that the decline in accuracy is directly linked to the transition abruptness determined by deme connectivity. Therefore, SMC-based algorithms are expected to perform better when applied to more isolated demes, where transitions between the *IICR*_*Sim*_ values in the ancient times and the recent times are smoother.

We also observed that the error of PSMC in FIM models increases with *M* for small *d*, whereas for large *d*, the error is greatest at low *M* . This pattern reflects both the general difficulty of SMC-based methods in recovering pronounced demographic changes in a few generations (Sellinger et al., 2021) (i.e. the degree of abruptness of the transition phase) and the specific limitation of PSMC in resolving recent times (Patton et al., 2019; Terhorst et al., 2017), which is observed in our simulations even after excluding the first 50 generations. The expected shape of the *IICR*_*Sim*_ contextualizes these biases. When *d* is large, the difference between IICR values at ancient and recent times becomes more pronounced, which amplifies errors in PSMC inference. At moderate migration rates (*M* ), the transition between the collecting and scattering phases is shifted toward recent times (Figure 1), where PSMC is known to perform poorly (Figure 3). In contrast, when the number of demes (*d*) is moderate and migration rate (*M* ) is low, the difference between the collecting and scattering phases is smaller and the transition occurs at more ancient times. In these scenarios, PSMC is able to accurately capture the *IICR*_*Sim*_ during both the transition period and the recent past (scattering phase) (Figure 3).

In the case of SMC++, beyond the general increase in error with transition abruptness —and thus with deme connectivity— and the reduced accuracy in scenarios with a large number of demes, we also observed higher error for low values of N. This suggests that the algorithm struggles to recover very small recent *IICR*_*Sim*_ values, likely as a consequence of the reduced number of coalescent and recombination events in small demes, which limits the information available for inference and lowers the accuracy of SMC++.

Our results indicate that the parameters of migration rate (*M* ) and number of demes (*d*) are crucial factors shaping the accuracy of coalescent-based inferences. Both algorithms lose accuracy in highly fragmented scenarios (large *d*) and in less isolated demes, but their response to migration (*M* ) differs: PSMC performs better at low migration rates, while SMC++ is more accurate at high migration. This complementary behavior suggests that the two algorithms can be jointly informative for estimating the IICR. In the simple equilibrium structured scenario we tested here, the challenging part of the historical demography reconstruction seems to lie in the transition between the collecting and the scattering phase. However, it is possible that in more complex scenarios with changes in deme size and/or connectivity through time he two algorithms may fail to detect such changes and may behave differently, something which remains to be explored. One of the key findings of this study is that both algorithms exhibit systematic biases inducing an overestimation of the IICR at specific time points, that could be misinterpreted as a demographic expansion if panmixia is assumed.

#### PSMC bias

Looking forward in time, we detected a spurious expansion at the beginning of the transition phase under specific parameters’ combination. The peak is followed by a population bottleneck after which a plateau is reached until recent times, with an *IICR*_*PSMC*_ higher than the simulated one. This pattern recurrently appears in real data in many studies (Hilgers et al., 2025) but here we provide some hints to when this may occur. Indeed, we investigated the probability of finding the spurious peak as a function of demographic parameters and found that it is highest at moderate values of *M* and with large numbers of demes (*d*). The bias was stronger in FIM scenarios than in 2D-SST scenarios, where it appeared only in the central deme and was absent in the corner deme. Thus, we find that the bias is linked to more abrupt transition periods, which occur when the focal deme is more connected to the rest of the metapopulation. Consistently, we also found that the size of the bias increases with the number of connected demes. It has been recently proposed to get rid of this bias by playing with the atomic time vector of the PSMC to increase the number of recent time intervals (Hilgers et al., 2025). However, the ultimate goal of applying SMC methods to genomic data is to uncover the complex demographic history of the population, i.e., to understand the evolutionary process that gave origin to the observed IICR. While it could prove difficult to find the right atomic vector in real datasets where the true IICR is unknown, our results provide a clear link between the presence of this peak and the underlying population structure. This observation is of practical utility to put forward evolutionary hypotheses to be tested with model based approaches, in which unsampled populations could be added to the inferential process using SFS (Excoffier et al., 2013, 2021) or LD based (Ragsdale, 2022) statistics.

#### SMC++ bias

Looking forward in time, the spurious expansion was always detected at the end of the transition period (Figure 7). The start of the expansion in our simulated framework was always observed between 1200 and 250 generations before present, a time frame that would not be discarded in empirical studies for being too recent. In other words, the spurious signal would be interpreted as a real demographic expansion under the panmixia hypothesis.

The probability of detecting this spurious expansion increases with low migration rates and a large number of demes. These conditions correspond to metapopulations with strong population structure, where the waiting times for coalescence events are longer as lineages need more time to end up in the same deme. Intuitively, the long time spent in different demes of the surviving lineages determines an accumulation of differentiation: once the lineages end up in the same deme they will suddenly increase the observed heterozigosity. To cope with this excess of diversity, SMC++ reconstructs a spurious recent expansion. This interpretation is further supported by the strong positive association we observed between the probability of detecting the artificial expansion and *Fst*. Structured scenarios with *Fst* values above 0.27 show more than a 50% probability of producing this false signal. *Fst* is an empirical measure that can be easily obtained from real data, providing a practical way to anticipate when SMC++ inference is most likely to be biased.

The spurious expansion detected by the SMC++ is similar to the bias recently observed when applying GONE (Novo et al., 2023; Santiago et al., 2020) to structured scenarios with low migration rates. GONE models the observed LD statistics and focuses on the last tens of generations, a time scale not considered in our study due to the intrinsic limits of both PSMC and SMC++. The convergence between our results and the bias observed with GONE suggests that the underlying mechanism behind the spurious increase is likely a general limitation of demographic inference methods assuming panmixia when reconstructing recent histories in structured populations.

### Implications for empirical studies and further research

Our study highlights the importance of explicitly interpreting the inferred IICR under structured models. We demonstrate that the performance of PSMC and SMC++ is heavily influenced by the demographic parameters of the underlying scenarios. Two recurrent biases consistently emerge, both of which related to specific parameters’ combination and suggesting population expansion if interpreted under the panmixia hypothesis.

Rather than dismissing them as an artifact, these biases can be seen as heuristic predictors of population structure, given that: (i) the probability of retrieving a recent spurious expansion with the SMC++ is positively correlated to the *Fst*; and (ii) the PSMC bias is related to the degree of connectivity of the sampled deme and the total number of demes of the metapopulation. Interpreting these biases in the light of hidden population structure in empirical datasets can provide useful information about the evolutionary history of a species, particularly if corroborated by independent metrics and data.

Recent theoretical advances show that the SMC framework could be extended to predict the IICR trajectory in a two-population admixture model and compute the likelihood that a dataset originates from structured versus panmictic models (Cousins et al., 2025). While these approaches offer a promising path forward, their ability to be scaled up to more complex scenarios, such as constant migration among many subpopulations, isolation by distance, range expansions or time-varying connectivity, remains an open challenge.. Future research should focus on quantitative approaches to discriminate between complex population structure and panmixia and to extend the SMC framework to more complex and realistic demographic scenarios. At the moment, SMC based models should be used as powerful descriptors of the distribution of the coalescence rate through time. Together with their identified biases, they can provide essential insights into the evolutionary history of natural species and can help to put forward specific hypotheses about their historical demography to be tested with complex model based approaches.

## Acknowledgements

AN acknowledges support from the European Union’s Horizon 2020 research and innovation programme under the Marie Skłodowska-Curie grant agreement No 945304 – Cofund AI4theSciences hosted by Université PSL

## Data Accesibility and Benefit-Sharing

All code to perform simulations and analysis can be found in the following Github repository: https://github.com/albanieto/structured_IICR_SMC.git

## Author Contributions

AN conducted analysis, study design and wrote manuscript. OL study design and wrote manuscript. SM study design and wrote manuscript.

## Notes

### Competing Interest Statement

The authors have declared no competing interest.

