## Supplementary Material for "Performance of Sequential Markovian Coalescence Methods when Populations are Structured"


### Contents


1. Obtaining the IICR 2
   1.1 $IICR_{Sim}$ 2
   1.2 $IICR_{Inf}$ 3
      1.2.1 $IICR_{PSMC}$ 3
      1.2.2 $IICR_{SMC++}$ 4
2. Comparison of $IICR_{Sim}$ and $IICR_{Inf}$ 5
3. Supplementary Figures 6
   Sup. Figure 1. SOM in $IICR_{Sim}$ separating FIM models
   Sup. Figure 2. SOM in $IICR_{Sim}$ separating 2D-SST models
   Sup. Figure 3. PSMC inference overall error in 2D-SST
   Sup. Figure 4. SMC++ inference overall error in 2D-SST
   Sup. Figure 5. Probability of artifacts in PSMC and SMC++ in FIM
   Sup. Figure 6. Probability of artificial expansion in SMC++ vs Fst

### 1 Obtaining the IICR

For each scenario of FIM and 2D-SST (corner and central sampling), we obtained the $IICR_{Sim}$ and the $IICR_{Inf}$. The $IICR_{Sim}$ is the expected trajectory of the IICR of a simulation and the $IICR_{Inf}$ was inferred with two inference algorithms, PSMC and SMC++.

#### 1.1 $IICR_{Sim}$

To calculate the $IICR_{Sim}$, we simulated $10^7$ $T_2$ extracted from the target deme under each scenario. The coalescence times were collected using the flag $-recordMRCA$ in *fastsimcoal2* (Excoffier et al., 2013). To obtain the cumulative density function ($F_2$) of $T_2$, we defined a vector $v$ of $n = 200$ logarithmically spaced time points between 50 and $5 \times 10^4$ generations for FIM models. For models from groups B, 2D-SST sampling from the central and the corner deme, and C, pairs of FIM and 2D-SST with equivalent demographic parameters, the number of time points was increased to $n = 400$, covering a broader range from 50 to $10^5$ generations. This increase was necessary to capture the variation in $IICR_{Sim}$, as the duration of the collecting phase —directly proportional to $Nd$, and inversely proportional to $M$ (Wakeley, 1999)— is longer in 2D-SST models. We used the vector v to create a histogram of the distribution of coalescent times $T_2$ and calculated the $IICR_{Sim}$ following Equation 1 from the main text as in Mazet et al. (2016).

$$IICR(t_i) = \frac{1 - F_{T_2}(t_i)}{f_{T_2}(t_i)}$$

#### 1.2 $IICR_{Inf}$

##### 1.2.1 $IICR_{PSMC}$

We simulated five genomic regions of 100Mb each for each of the 20 haploids, randomly coupled to obtain ten diploid individuals. Since the convergence of SMC-based algorithms is known to depend on the ratio of the mutation rate to the recombination rate (Sellinger et al., 2021), we fixed both parameters to $1 \times 10^{-8}$ per site per generation, an order of magnitude consistent with many empirical studies (Hudson, 2002). For PSMC analysis, we considered the default parameters of bin size $-s100$, maximum time interval $-t15$, and time vector $-p4+25*2+4+6$ (Li and Durbin, 2011). While adjusting these parameters can potentially enhance PSMC performance (Hilgers et al., 2025; Mather et al., 2020; Peede et al., 2025), we considered the defaults as these are the ones usually applied to species for which we lack prior knowledge (Mattle-Greminger et al., 2018; Palkopoulou et al., 2015; Patton et al., 2019; Qi et al., 2023), and to avoid introducing bias through model-specific tuning. We ran PSMC for each of the 10 haploid pairs sampled from the deme. Subsequently, we interpolated the time points in vector $v$ for each resulting inference and calculated the arithmetic mean to derive the $IICR_{PSMC}$.

##### 1.2.2 $IICR_{SMC++}$

We followed a similar approach with SMC++, where we obtained 10 independent inferences by setting as *distinguished* individual each of the locally sampled pairs at a time, and as *undistinguished* individuals the other nine. We set up the starting time as 0 and the number of timepoints for the estimation to $1 \times 10^5$ generations. Then, we interpolated the timepoints in the vector $v$ from each of the 10 individual's inferences (specified in the final output of an SMC++ run as $y$) and computed the arithmetic mean to obtain $IICR_{SMC++}$ of each simulation.

#### 2 Comparison of $IICR_{Sim}$ and $IICR_{Inf}$

For each scenario (and sampling in the case of 2D-SST), we compared the expected IICR in a simulation, the expected IICR in a simulation ($IICR_{Sim}$), with the $IICR_{Inf}$ obtained with each of the two inference algorithms, the $IICR_{PSMC}$ and the $IICR_{SMC++}$. To do so, we calculated the root of the mean squared error ($RMSE$):

$$\mathrm{RMSE} = \sqrt{\frac{1}{n}\sum_{i=1}^{n}(\mathrm{IICR}_{\mathrm{Sim},i} - \mathrm{IICR}_{\mathrm{Inf},i})^2}$$

Here $i$ represents each of the timepoints in vector $v$ and $n$ the number of timepoints in vector $v$.

#### 3 Supplementary Figures

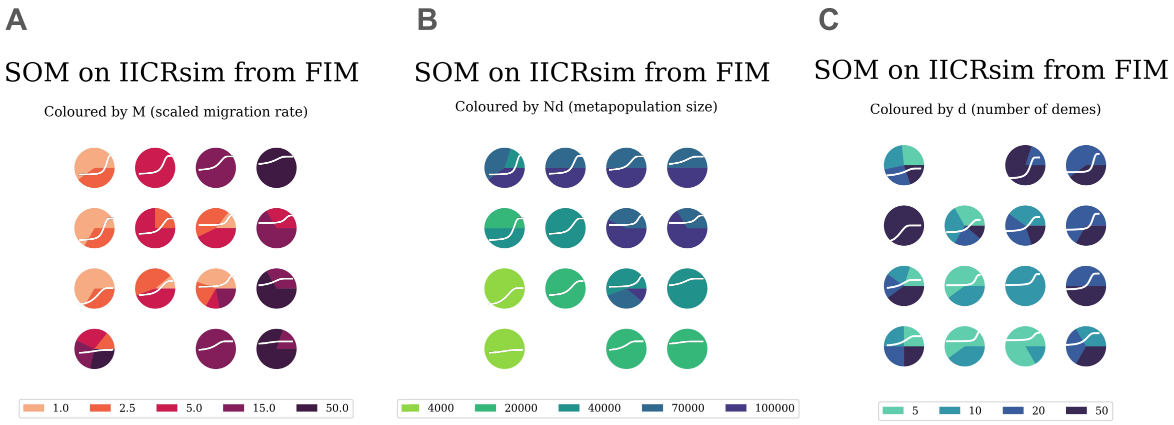


Figure S S1: **SOM in $IICR_{Sim}$ separating FIM models by parameters** Self organizing Map of 4x4 neurons. $IICR_{Sim}$ curves are categorized depending on the value of a specific demographic parameter: $M$ (**A**), $Nd$ (**B**) and $d$ (**C**). In map **C**, scenarios with $M = 50$ are not considered. Pie charts represent the proportion of $IICR_{Sim}$ with a specific parameter value placed in a specific neuron. The value of the demographic parameter is designed by a scale that goes from light –for smaller values– to dark colours –for larger values–. The winner $IICR_{Sim}$ curve is placed on the top of each neuron in white.

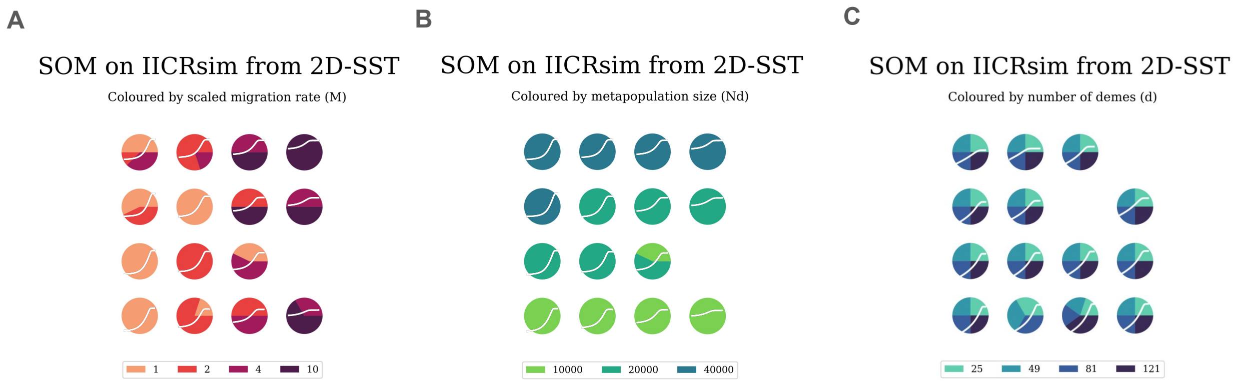


Figure S S2: **SOM in $IICR_{Sim}$ separating 2D-SST models by parameters** Self organizing Map of 4x4 neurons. $IICR_{Sim}$ curves are categorized depending on the value of a specific demographic parameter: $M$ (**A**), $Nd$ (**B**), $d$ (**C**). Pie charts represent the proportion of $IICR_{Sim}$ with a specific parameter value placed in a specific neuron. The value of the demographic parameter is designed by a scale that goes from light –for smaller values– to dark colours –for larger values–. The winner $IICR_{Sim}$ curve is placed on the top of each neuron in white.

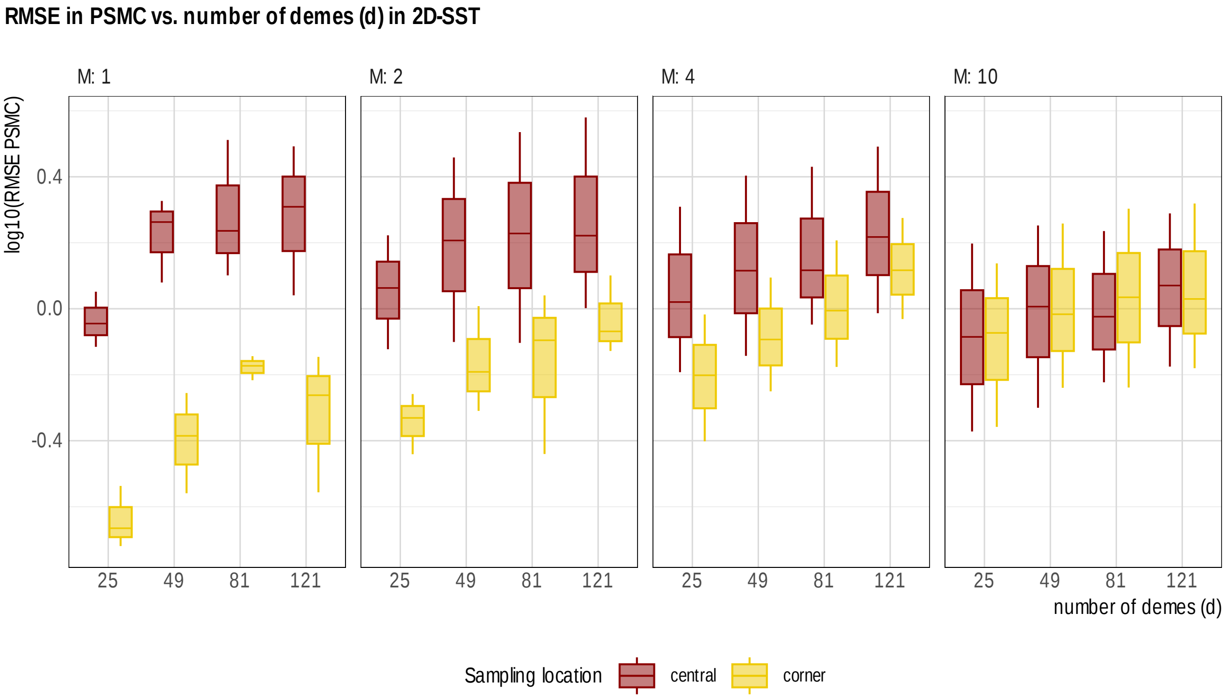


Figure S S3: **PSMC inference overall error in 2D-SST** Models are separated in panels depending on the value of scaled migration rate ($M$) (indicated above each graph). Vertical axis corresponds to the decimal logarithm of the $RMSE_{PSMC}$ and horizontal axis represents the values of number of demes ($d$). Colors represent the sampling from the corner deme (yellow) or the central deme (red).

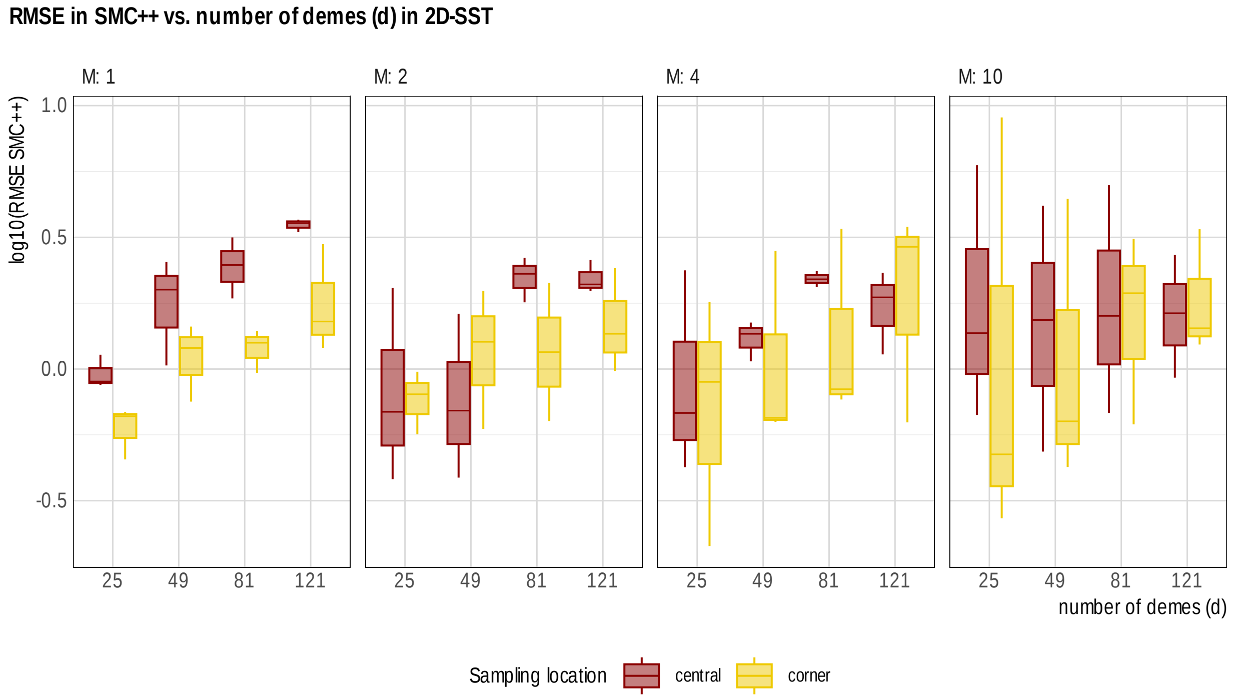


Figure S S4: **SMC++ inference overall error in 2D-SST** Models are separated in panels depending on the value of scaled migration rate ($M$) (indicated above each graph). Vertical axis corresponds to the decimal logarithm of the $RMSE_{SMC++}$ and horizontal axis represents the values of number of demes ($d$). Colors represent the sampling from the corner deme (yellow) or the central deme (red).

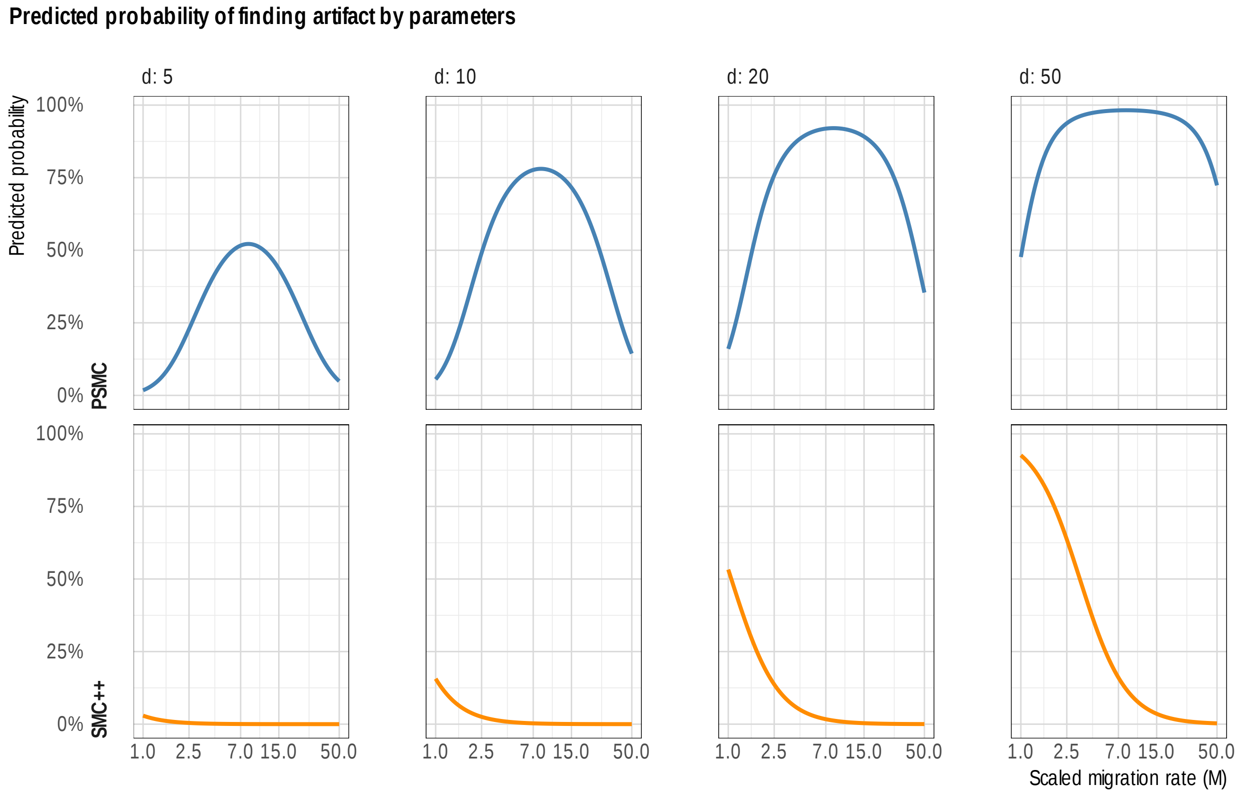


Figure S S5: **Probability of finding the artifacts in PSMC and SMC++ given demographic parameters in FIM** Probability of finding the artificial expansion in PSMC (blue, upper panel) and the artificial recent recovery in SMC++ (orange, lower panel) in FIM in the case of $N = 2000$. Subplots are separated by the value of the number of demes considered ($d$). Vertical axis represents the probability of finding the artifact; horizontal axis represents the value of $M$.

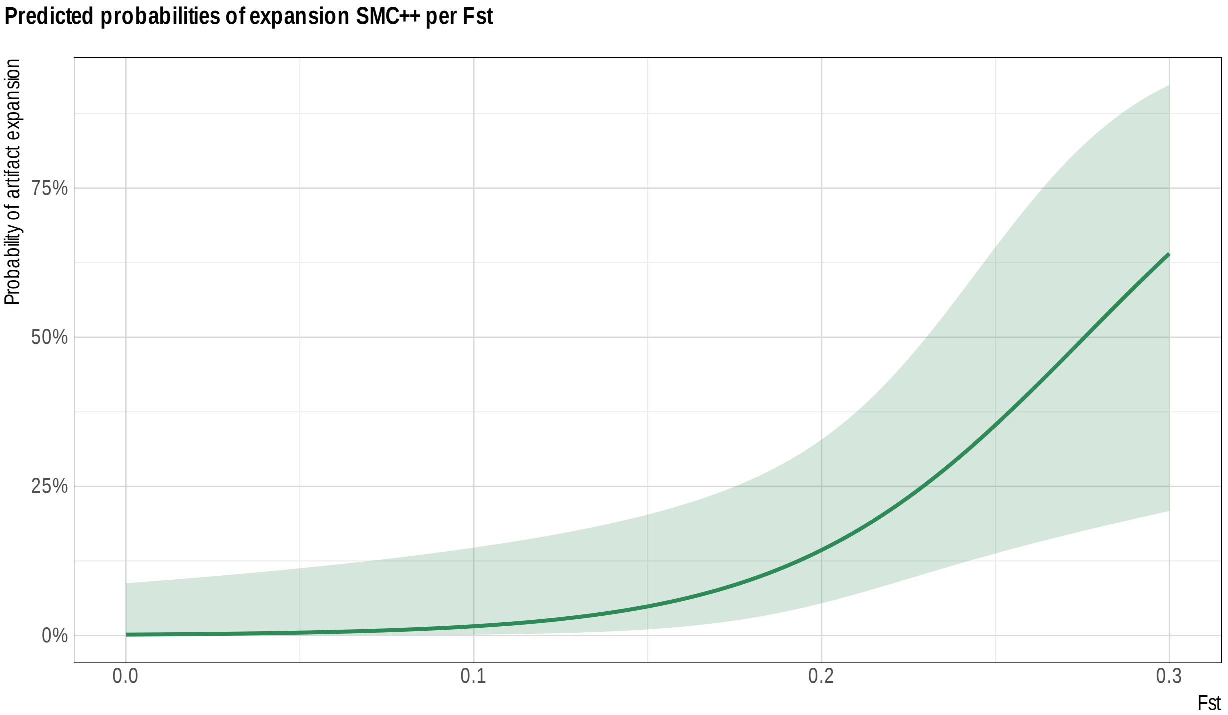


Figure S S6: **Probability of finding artificial expansion in SMC++ in FIM models depending on Fst** Vertical axis represents the probability of finding the artificial recent expansion and horizontal axis the Fst value. Green line represents the mean modeled probability at each Fst value and the area corresponds to the 95% confidence interval.

David Peede, Trevor Cousins, Arun Durvasula, Anastasia Ignatieva, Toby G. L. Kovacs, Alba Nieto, Emily E. Puckett, and Elizabeth T. Chevy. Not Just $N_e$ $N_e$-more: New Applications for SMC from Ecology to Phylogenies, 2025.

Xiao-Guang Qi, Jinwei Wu, Lan Zhao, Lu Wang, Xuanmin Guang, Paul A. Garber, Christopher Opie, Yuan Yuan, Runjie Diao, Gang Li, Kun Wang, Ruliang Pan, Weihong Ji, Hailu Sun, Zhi-Pang Huang, Chunzhong Xu, Arief B. Witarto, Rui Jia, Chi Zhang, Cheng Deng, Qiang Qiu, Guojie Zhang, Cyril C. Grueter, Dongdong Wu, and Baoguo Li. Adaptations to a cold climate promoted social evolution in Asian colobine primates. *Science*, 380(6648): eabl8621, June 2023. ISSN 0036-8075, 1095-9203. doi: 10.1126/science.abl8621.

Thibaut Paul Patrick Sellinger, Diala Abu-Awad, and Aurélien Tellier. Limits and convergence properties of the sequentially Markovian coalescent. *Molecular Ecology Resources*, 21(7): 2231–2248, 2021. ISSN 1755-0998. doi: 10.1111/1755-0998.13416.

John Wakeley. Nonequilibrium Migration in Human History. *Genetics*, 153(4):1863–1871, December 1999. ISSN 1943-2631. doi: 10.1093/genetics/153.4.1863.
